# *GIGANTEA* is required for robust circadian rhythms in wheat

**DOI:** 10.1101/2024.04.19.590265

**Authors:** Laura J. Taylor, Gareth Steed, Gabriela Pingarron-Cardenas, Lukas Wittern, Matthew A. Hannah, Alex A. R. Webb

## Abstract

GIGANTEA (GI) is a plant-specific protein that functions in many physiological processes and signalling networks. In Arabidopsis, GI has a central role in circadian oscillators regulating the abundance of ZEITLUPE and TIMING OF CAB EXPRESSION 1 proteins and is essential for photoperiodic regulation of flowering. We have investigated how ortholgues of this component of Arabidopsis circadian oscillators contributes to circadian rhythms and yield traits, including heading (flowering) in wheat. We find that *GI* is a core component of wheat circadian oscillators that is necessary to maintain robust oscillations in chlorophyll fluorescence and circadian oscillator transcript abundance. Predicted lack of functional *GI* results in later flowering in wheat in both long days and short days in controlled environment conditions. Our results support and extend previous work which suggests that the pathways by which photoperiodism regulates flowering are not fully conserved between Arabidopsis and wheat. Understanding the molecular basis for photoperiodism in wheat is important for breeders looking to manipulate flowering time and develop new elite, high yielding cultivars.

**Highlight:** *GIGANTEA* is required for robust circadian oscillations in wheat and regulates heading, most likely through a *PHOTOPERIOD-1*-dependent pathway.

## Introduction

Circadian oscillators are endogenous timing mechanisms that act to time internal processes to daily and seasonal cycles in their environment. In the model plant Arabidopsis, well defined transcription-translation feedback loops result in sequential expression of circadian oscillator genes across the 24-hour cycle. At dawn, *CIRCADIAN CLOCK ASSOCIATED 1* (*CCA1*) and *LONG ELONGATED HYPOCOTYL* (*LHY*) are expressed and repress the day phased *PSEUDO RESPONSE REGULATOR* (*PRR*) genes (*PRR7*, *PRR9* and *TIMING OF CAB EXPRESSION 1*, *TOC1*) (Harmer *et al*., 2000; Adams *et al*., 2015). REVILLE (RVE)/ LIGHT NIGHT INDUCIBLE AND CLOCK REGULATED 1 (LNK1) complex promotes *PRR* expression (Rawat *et al*., 2011; Xie *et al*., 2014), which feeds back to repress *CCA1*/*LHY* (Nakamichi *et al*., 2010). The evening complex (EARLY FLOWERING 3/EARLY FLOWERING 4/LUX ARRYTHMO) represses *PRR* expression at dusk (Dixon *et al*., 2011; Helfer *et al*., 2011; Nusinow *et al*., 2011). PRR action is further restricted to the light by ZEITLUPE (ZTL)-mediated degradation of PRR proteins in the dark (Cha *et al*., 2017; Lee *et al*., 2019). Thus, *CCA1*/*LHY* repression is lifted towards the new dawn and the cycle repeats. In Arabidopsis, proteins that might function as scaffolds have profound effects on the circadian oscillator. For example, members of the WD40 repeat family of scaffold proteins are essential for circadian rhythms (Airoldi *et al*., 2019). GIGANTEA (GI) is another potential scaffold protein that is important in circadian timing. GI regulates ZTL stability and activity through light-dependent binding and affects deubiquitination and also translational activity (Lee *et al*., 2019). Mutations in *GI* have allele-specific effects on Arabidopsis circadian oscillators but are mostly considered to result in faster running circadian oscillators and therefore shorter circadian periods in constant conditions (Hsu and Harmer, 2014). The architecture of circadian oscillators is broadly conserved across the plant kingdom, with alleles of circadian oscillator genes selected during domestication and in breeding programs to enhance agriculturally important crop traits, such as flowering time (McClung, 2021; Steed *et al*., 2021). However, we have found in wheat that transcripts of *ELF3*, which encodes a putative scaffold protein, peak at dawn, rather than dusk as in Arabidopsis (Wittern *et al.,* 2023). Furthermore, wheat ELF3 proteins are unstable in the light (Alvarez *et al.,* 2023). These data suggest that there are differences in circadian oscillator structure and function in wheat compared to Arabidopsis and warrant further investigation.

In Arabidopsis, mutations in *GI* also have profound effects on the photoperiodic flowering pathway due to both its function in regulating circadian oscillators and more direct roles in regulating the floral induction mechanisms (Fowler *et al*., 1999). In the external coincidence model of photoperiodism, flowering is induced when external light is coincident with an appropriate phase of the circadian rhythm (Bunning, 1960; Pittendrigh and Minis, 1964). GI connects circadian oscillators and the photoperiodic flowering pathway, and along with FLAVIN BINDING KELCH REPEAT 1 (FKF1) and CONSTANS (CO) are the molecular components which determine the light-sensitive and -insensitive phase of the circadian cycle for the regulation of flowering (Suárez-López *et al*., 2001; Valverde *et al*., 2004; Sawa *et al*., 2007). In long photoperiods, *GI* and *FKF1* peak expression coincides in the late afternoon, forming a complex which promotes *CO* expression. In Arabidopsis, if light is coincident with the accumulation of *CO* transcripts, CO protein is stabilised to pass the threshold necessary to trigger *FLOWERING LOCUS T* (*FT*) expression (Imaizumi *et al*., 2003; Sawa *et al*., 2007; Fornara *et al*., 2009). Arabidopsis FT is a mobile signal which travels from the leaf to the apical meristem to trigger floral development (Corbesier *et al*., 2007). In short photoperiods, *FKF1* and *GI* maximal expression do not coincide, hence *CO* repression is not lifted during the day and maximal CO accumulation occurs at night, outside of the light sensitive phase of the rhythm (Sawa *et al*., 2007). Therefore, GI mutants such as *gi-1* (premature stop coding resulting in the loss of 171 of the C-terminal domain*), gi-2* (causing the deletion of all but 142 amino acids of the N-terminal domain and addition of the 16 new amino acids in the C-terminal domain) and T-DNA insertion line (*gi-11,* deletion in the 5’ half of *GI* caused by the insertion of the T-DNA) are late flowering in long photoperiods but have no, or minor delays to flowering when cultivated in short photoperiods (Fowler *et al*., 1999; Park *et al*., 1999).

In wheat photoperiodic-dependent regulation of flowering is heavily dependent on *Photoperiod 1* (*PPD-1*), an orthologue of Arabidopsis *PRR3* or *PRR7* (Turner *et al*., 2005; Beales *et al*., 2007; Wilhelm *et al*., 2009). *PPD-1* induces *FT1* expression in leaves activating the transition to the reproductive stage (Alvarez *et al*., 2023). Analyses of the tetraploid wheat loss of function Target Induced Local Lesions In Genome (TILLING) lines containing stop codon and splice site mutations have demonstrated that *PPD-1* and *CO1/CO2* act in separate but highly connected flowering pathways (Shaw *et al*., 2020). In tetraploid wheat, light activation of *PPD-1*, mediated by phytochromes PHYB and PHYC, is facilitated by ELF3, which represses the expression of *PPD-1* by directly binding to its promoter (Alvarez *et al*., 2023). *ELF3* controls the abundance of *PPD-1* to regulate flowering somewhat independent of its role in circadian oscillators in tetraploid wheat (Wittern *et al*., 2023). Tetraploid wheat plants carrying a deletion in the promoter of the A homoeologue of *PPD-1* (*Ppd-1Aa)* are photoperiod-insensitive (PI) and are early heading under short photoperiods. In contrast, plants that carry the *Ppd-1Ab* allele are photoperiod-sensitive heading later under short photoperiods (Alvarez *et al*., 2023).

In wheat *GI* regulates flowering, though the mechanisms are not yet resolved. Association studies using wheat lines from diverse geographical regions found that polymorphisms in *GI* are associated with earlier flowering (Rousset *et al*., 2011). In both wheat and barley, *GI* expression levels peak in the afternoon in long and short photoperiods (Dunford *et al*., 2005; Zhao *et al*., 2005; Rees *et al*., 2022). Wheat *GI* rescues the late flowering phenotype in Arabidopsis *gi-2* with over-expression of the wheat *GI* in Arabidopsis resulting in earlier flowering (Zhao *et al*., 2005). *GI* loss-of-function mutants in a photoperiod-sensitive wheat background have delayed heading dates in both long and short photoperiods. The stronger effects of *GI* on heading date in long photoperiods required *PPD-1* or *ELF3,* suggesting function in a common pathway (Li *et al*., 2024).

To investigate the relationship between the function of the circadian oscillator in the regulation of heading, we isolated wheat TILLING lines which carry predicted non-functional alleles of the Durum wheat orthologues of Arabidopsis *GI*. We demonstrate that *GI* is required for robust circadian oscillations in Durum wheat. We then investigated whether *GI* functions in the Durum wheat photoperiodic flowering pathway independent of normal *PPD-1* function. Finally, we investigated if lack of functional *GI* affects other aspects related to yield traits.

## Materials and Methods

### Isolation of *GI* TILLING mutant lines in tetraploid wheat

To investigate the role of *GI* in wheat, we isolated TILLING mutants in the tetraploid wheat variety *T. turgidum cv. Kronos*. Two TILLING lines with premature stop codons within the A *GI* homoeologues (*GI-A*, TraesCS3A02G116300, Kronos2019) and B *GI* homoeologues (*GI-B*, TraesCS3B02G135400, Kronos2205) genes were obtained from the Germplasm Resource Unit (GRU), Norwich UK (Figure 1). BLAST search of the sequenced Kronos TILLING lines for *TtGI* identified pairs of candidate lines with deleterious mutations in each of the subgenomes for *TtGI*, which were obtained directly from Dr. Cristobal Uauy (John Innes Centre, Norwich). Further candidates from tetraploid Kronos were identified using a pre-publication version of the http://www.wheat-tilling.com website (Krasileva *et al*., 2017). Kronos2019 contains a C to T SNP 6606 nucleotide base pairs (bp) from the start of the *GI*-*A3* transcription start site (TSS) (position 3A:84190982 IWGSC Refseq V1.1). This SNP is situated within the 12^th^ exon of the first splice variant and the fourth exon of the second splice variant and is predicted to result in the deletion of 356 amino acids (Fig. 1A). Kronos2205 contains a G to A SNP 6348 bp downstream of the *GI-B3* TSS (position 3B:117929486 IWGSC Refseq V1.1). This SNP is situated within the 12^th^ exon of the first two splice variants and the 11^th^ exon of the third splice variant and is predicted to result in the deletion of 403 amino acids (Fig. 1B). Abundance of *TtGI* at ZT12 and ZT15 is not significantly different between the wild type genotypes, single and double mutants, with both homologous expressing at the same level (Supplementary Fig. S1).

**Fig. 1.**
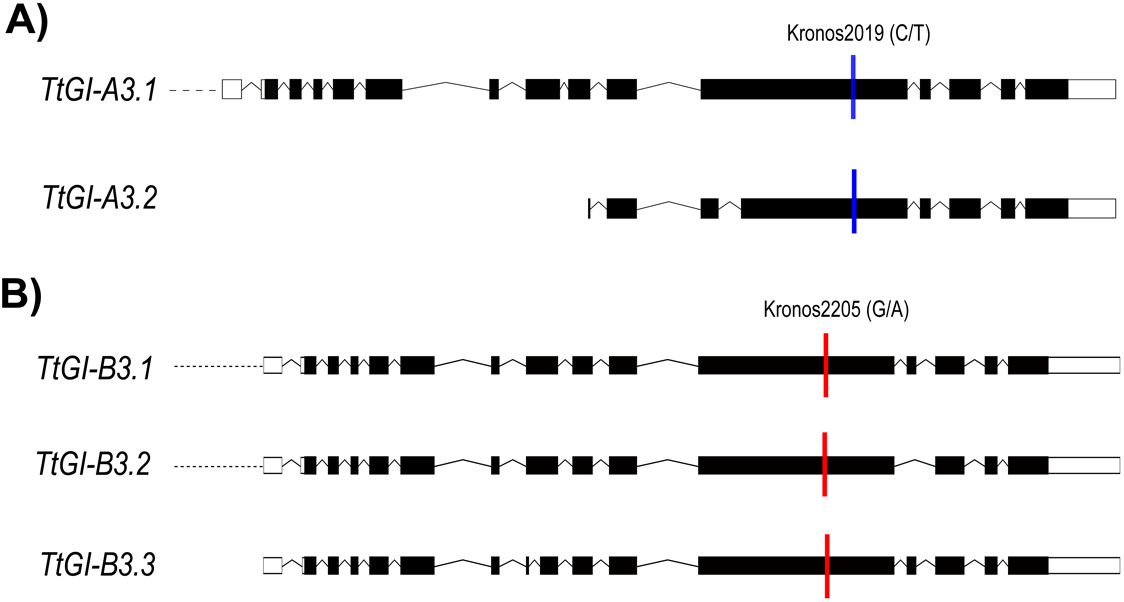
Location of TILLING mutations in the *GI-A3* and *GI-B3* homeologues. Location of premature stop codons (coloured bars) in (A) *GI-A3* and (B) *GI-B3* homoeologues (showing their respective splice variants, genetic alterations resulting in an altered coding sequence). Bars represent exons with black and white showing those that are translated and untranslated, respectively. Kronos2019 and Kronos2205 are the original TILLING lines obtained from the GRU. Schematic was adapted from EnsemblPlants database.

Kronos TILLING lines were backcrossed four times to Kronos (Supplementary Fig. S2). The BC4 F1 heterozygous plants were self-crossed to produce BC4 F2 seed. BC4 F2 seeds were genotyped, and seeds retained from plants which were homozygous for the Kronos2019 or Kronos2205 mutations. The homozygous Kronos2019 (*Ttgi-A3)* and Kronos2205 (*Ttgi-B3)* lines were crossed, self-crossed and plants homozygous for both Kronos2019 and Kronos2205 mutations (*Ttgi-A3/gi-B3*) were selected. To control for possible background mutations, that are present in the TILLING populations, plants homozygous for both wild type alleles were selected as mutant sibling wild type (WT segregant). Kronos wild type line was selected as non-mutagenized wild type control.

### Genotyping

Genotyping was performed by PCR followed by Sanger sequencing conducted by Source Bioscience Ltd (UK). A Faststart Taq Polymerase Kit (Roche, UK) was used for PCR following the manufacturer’s instructions. PCR conditions were as follows: 4 min at 95°C; 40 cycles of 30 s at 95°C, 30 s at 68°C/69°C (*Ttgi-A3* and *Ttgi-B3* respectively), 60 s at 72°C; 10 min at 72°C. Primers used for the A homologue: forward 5’ GCATCATCCCTCTACCATTTGA 3’ and reverse 5’ GTACAAGCTTCACCGTCGA 3’. Primers used for the B homologue: forward 5’GTTGGTACCTCCTGGAACTACA 3’ and reverse 5’CATCCCATCTGTAGCACGAAGTA 3’.

### Plant Growth conditions

Seeds were sown directly into modular trays containing a 5:1 mix of Levington M2 potting compost pre-treated with Intercept 70W (0.02 gL^-1^, Bayer) and fine vermiculite. Seeds were stratified for 48 hours at 4°C and then moved into a growth cabinet (PGR14, Conviron) or room (Conviron) fitted with broad spectrum LED white light. If plants were to grow to maturity, seedlings were transplanted into 12 by 12 by 13 cm pots containing an identical soil mix after two weeks of growth. Plants were grown under 400 µmol m^-2^ s^-1^ PAR, 22°C white light/ 18°C dark in either long (16 hours light/ 8 hours dark) or short (8 hours light/ 16 hours dark) photoperiods.

### Chlorophyll fluorescence measurements

Chlorophyll fluorescence (CF) measurements were performed as outlined in Wittern *et al*., (2023). A five mm by five mm leaf fragment was placed into a well of a black 96 well plate (Greiner) containing 0.8% (w/v) bactoagar (BD), ½ MS (Duchefa Biochemie) and 0.5 uM 6-benzyl aminopurine (Sigma), adjusted to pH5.7 using 0.5 M KOH (Sigma).

CF imaging and processing was performed using a CFImager and accompanying software (Technologica Ltd., UK). CF images were captured using a Stingray F145B ASG camera (Allied Vison Technologies, UK) through a RG665 long pass filter to exclude blue light from the LEDs. The ‘continuous light’ protocol included the following steps: 40 minutes 100 µmol m^-2^s^-1^ of blue light, 800 ms 6172 µmol m^-2^s^-1^ saturating light pulse and 20 minutes of darkness. This protocol was repeated 120 times. The parameters Non-photochemical quenching (NPQ) and *F_v_/F_m_* (maximum potential quantum efficiency of Photosystem II, PSII) were used for circadian analysis. Relative Amplitude Error (RAE) calculation and period estimates were generated using Biodare2 (Zielinski *et al*., 2014).

### Reverse-transcription quantitative PCR

The *Ttgi-A3/gi-B3* and Kronos wild type lines were sown and grown in a growth room under long day conditions (16 h L: 8 h D; 250 µmol m^−2^ s ^−1^, 20 °C). *TtGI* expression was analysed from 14-day old plants at *TtGI* peak expression, ZT12 and ZT15. For analysis of circadian oscillator transcript abundance, samples were collected after 14 days of sowing from the first true leaf every 3 h for 96 h. During the first 24 h, sampling was done under long day conditions (16 h L: 8 h D; 250 µmol m^−2^ s ^−1^, 20 °C) and then the cabinet was switched to continuous white light (250 µmol m^-2^ s ^-1^) and constant temperature (20 °C). Flowering gene expression was analysed at the three-leaf growth stage (14 days after sowing) leaves and in GS39 growth stage (flag leaf blade all visible) under long day conditions (16 h L: 8 h D; 250 µmol m^−2^ s ^−1^, 20 °C). Sampling commenced at time 0 for 24 h every 3 h.

Total RNA was extracted from leaves using the RNeasy Plant Mini Kit (Qiagen, UK) with an on-column DNAse digest (Qiagen, UK), concentration and quality was determined using the Nanodrop ND-1000 (Thermofischer scientific). cDNA was synthesized from 500 ng RNA using the RevertAid First Strand cDNA synthesis kit (Thermo Scientific, UK). Three technical replicates of gene-specific products were amplified in 10 µl reactions using QuantiNova SYBR Green PCR Kit (Quiagen) on a CFX384 Touch Real-Time PCR detection system (Bio Rad). Transcript levels were determined relative to the expression of two housekeeping genes, *RP15*, *RPT5A* and Ta22845 as described in Wittern *et al.,* (2023). Primer sequences are shown in Table 1.

**Table 1.**
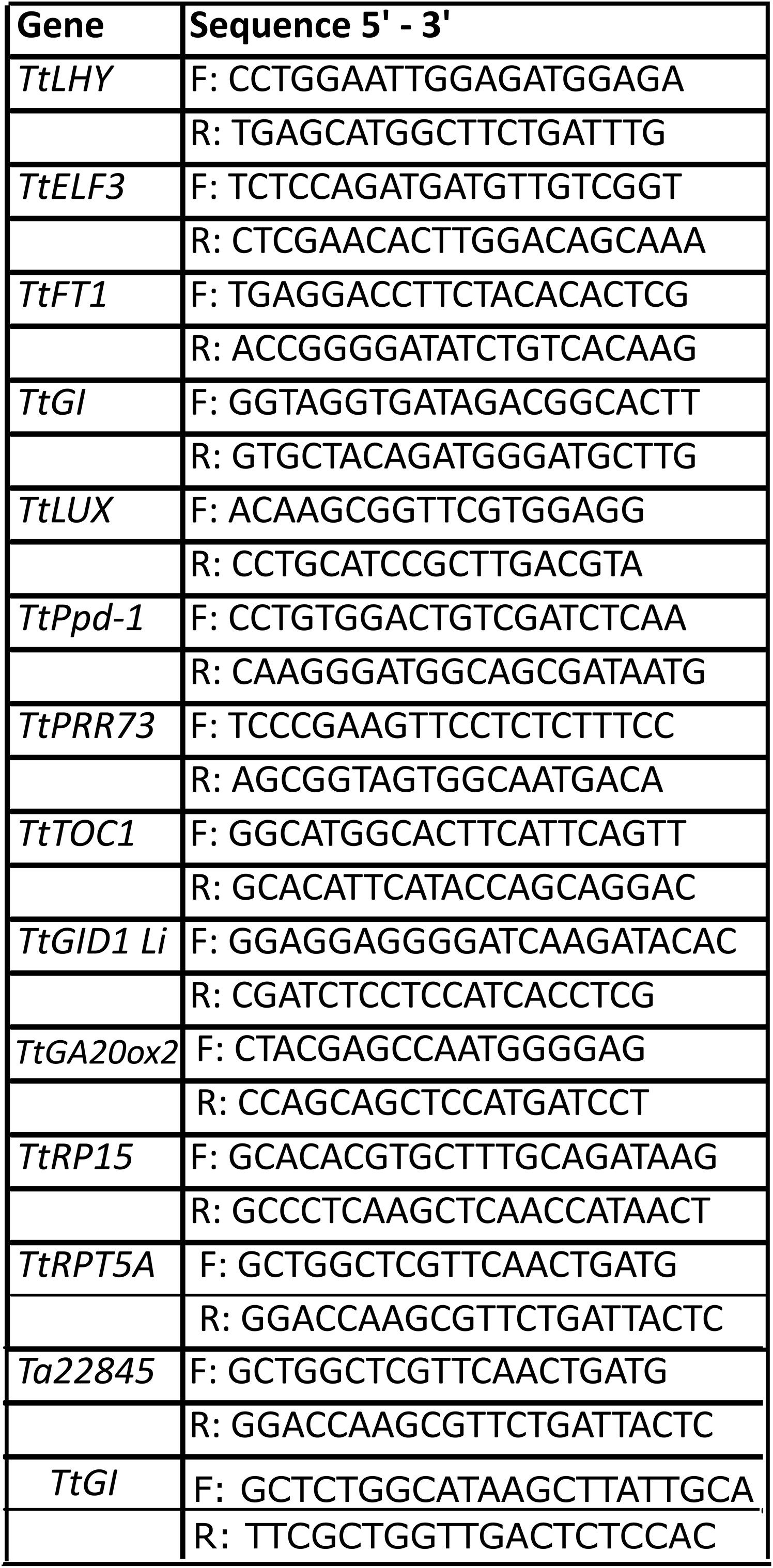
Primers used in this study.

### Plant phenotyping (laboratory)

Growth stage 55 (GS55), i.e., when half of the ear emerged above flag leave ligule (Zadoks *et al*., 1974) was recorded daily. The first tiller to reach GS55 (primary tiller) per plant was marked. Post senescence, the height of each plant was measured using a meter rule and the total number of tillers and number of productive tillers were counted. The primary head (head from primary tiller) was collected and the length, spikelet number, seed number and seed weight recorded. Seed number and weight were then determined for the whole plant. Seed number was counted by hand and seed weight measured using a balance.

### Field trial

Observational (1 m^2^) plots of Kronos, WT segregant, *Ttgi-A3* and *Ttgi-A3/gi-B3* lines were grown at National Institute of Agricultural Botany (NIAB, Cambridge, UK) experimental farm during the 2020 field season to provide preliminary data and bulk seed for a larger trial the following year. Environmental data is included in Supplementary information. On the 9^th^ of April 2021, Kronos, WT segregant, *Ttgi-A3* and *Ttgi-A3/gi-B3* yield plots (3.8m x 2m = 7.6 m^2^) were drilled at NIAB following a randomised block design (Supplementary Fig. S3). The trial was flanked by two rows of Paragon to separate the *GI* field experiment from other trials located at the site. *Ttgi-B3* was not included due to insufficient number of seed during the 2020 field season.

Five plants per plot were randomly tagged from the middle of each plot. The date each tagged plant reached GS55 was recorded. Post senescence, the height of each tagged plant was measured using a metre rule and the total number of tillers per tagged plant was counted. A representative head was taken from each tagged plant and the length, spikelet number, grain number and grain weight were recorded. Data was not recorded from Paragon control plants as the higher plant density and tillering meant that the tags were no longer visible within the Paragon plots. On the 4^th^ of September the plots were harvested and the weight of grain plus the percentage grain moisture for each plot was determined by the NIAB field team.

### Statistics

All statistical analysis was performed using R. For CF datasets, significant differences between the groups were tested for using an ANOVA followed by a post hoc Tukey test. Statistics for phenotypic data collected from plants grown in the laboratory and tagged plants from the field were as follows. Continuous variables (plant height, primary head length, weight of seed) were tested using an ANOVA followed by a Post hoc Tukey. All continuous discrete variables (days till GS55, tiller number, seed number, spikelet number) were tested using a Kruskal Wallis test followed by a post hoc Dunn test adjusted for multiple comparisons.

### Sequence analysis

*TaGI* genomic sequences and CDS were retrieved from the reference genome sequence (IWGSC RefSeqv1.1) and submitted as query sequences to local Basic Local Alignment Search Tool (BLAST) server to identify *GI* in wheat cultivar genomes released by the 10+ genomes project (Walkowiak *et al*., 2020) (Table S1). A reciprocal BLAST approach was used to confirm ambiguous results. Alignments were created using CLC Genomics between the BLAST results, the genomic BLAST query sequence (IWGSC RefSeqv1.1) the CDS and mRNA. SNPs and InDels within the CDS were manually annotated and counted.

## Results

### *TtGI* is required for robust circadian rhythms under constant light

To establish if *GI* contributes to the functioning of wheat circadian oscillators, we quantified circadian rhythms in chlorophyll *a* fluorescence from *Triticum turgidum cv. Kronos* by measuring the period and the relative amplitude error (measurement of the goodness of fit of the data to a cosine curve, RAE > 0.5 usually indicates a lack of circadian rhythms) (Fig. 2). Robust circadian oscillations of the chlorophyll *a* fluorescence derived parameter reporting non-photochemical quenching (NPQ) were measured in the recurrent parent and WT segregant lines (22.70 ± 0.27 hours, 23.30 ± 0.28 hours, respectively) (Fig. 2A and 2B). The measured periods were shorter than 24 h due to the high blue light intensity illumination used in our CF apparatus, which accelerates circadian oscillators and therefore shortens circadian period in constant conditions. Predicted loss of function of *Ttgi-B3* had little effect on circadian function (period 22.70 ± 0.22 hours, RAE 0.21 ± 0.01; Fig. 2F and 2G). Whereas plants in which there was a loss of functional *Ttgi-A3* had shorter period NPQ circadian rhythms (period 21.30 ± 0.14 hours, RAE 0.26 ± 0.01 Fig. 2F and 2G). The difference of period between single mutants was 1.40 hours. Mutation of both copies of *GI (Ttgi-A3/gi-B3*) resulted in loss of robust circadian rhythms demonstrated by an RAE of 0.59 ± 0.05 (Fig. 2F and 2G). The double mutant led to higher variation of plants that are rhythmic or arrhythmic. Here, 25% of the leaf fragments tested had a RAE < 0.5, indicating rhythmicity with a period of approximately 18h. The high variability of RAE values and shorter period is also seen in Arabidopsis *gi-3* mutant lines (Mizoguchi *et al*., 2005). Assessment of another chlorophyll *a* RAE parameter *F_v_/F_m_* (Supplementary Fig S4) resulted in the same conclusions that loss of *Ttgi-A3* shortens circadian period *(Ttgi-A3* period 21.71 ± 0.11 hours RAE = 0.27 ± 0.02, *Ttgi-B3* period 22.05 ± 0.11 hours, RAE *=* 0.26 ± 0.02), with a difference in period of 0.75 hours, and loss of both *Ttgi-A3* and *Ttgi-B3* abolishes robust circadian rhythms (RAE 0.52 ± 0.04; Supplementary Fig S4.G).

**Fig. 2.**
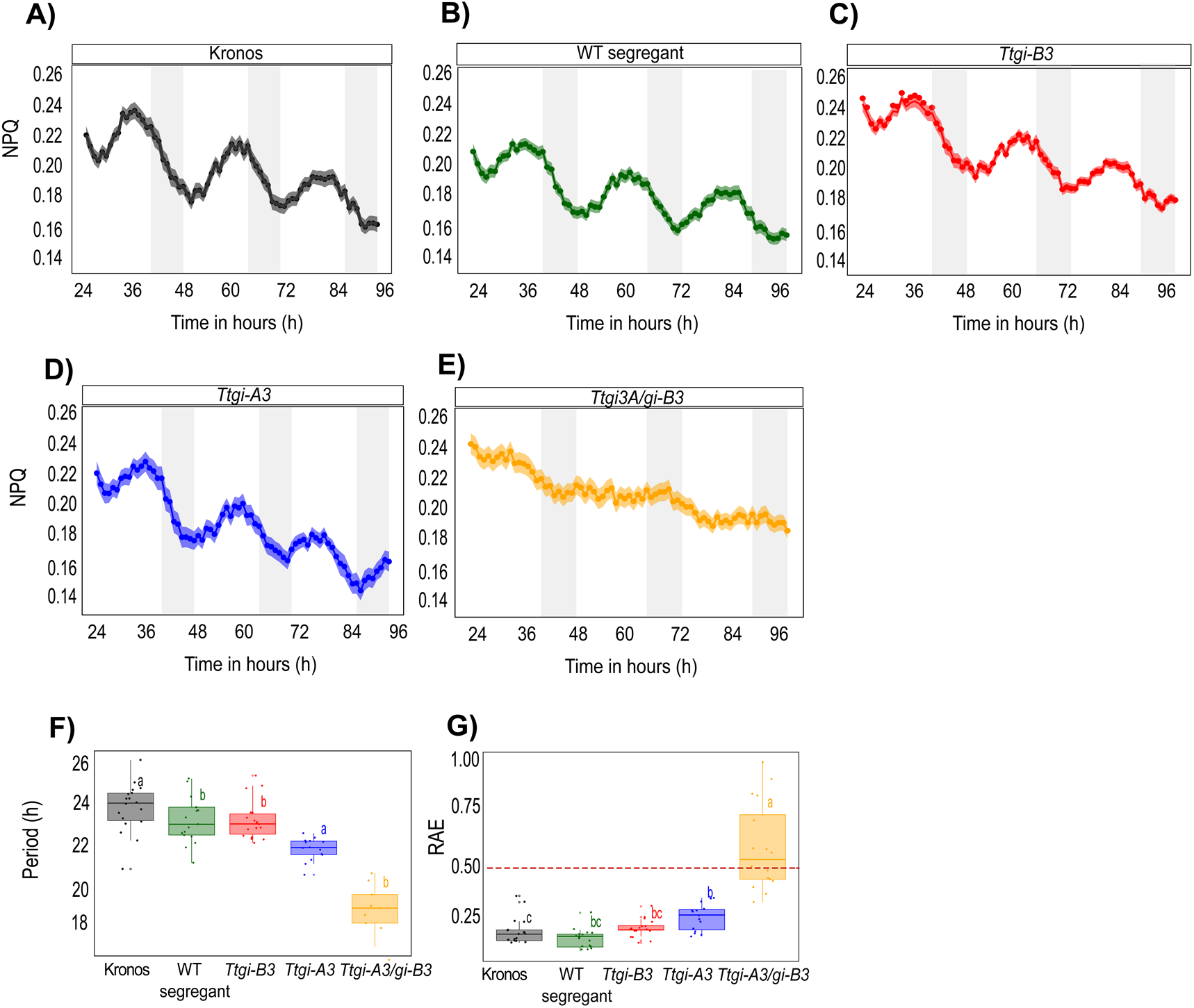
Functional *TtGI* is necessary for maintaining robust circadian rhythms in chlorophyll fluorescence. Mean of NPQ (± SEM, represented by the shaded ribbon) of Kronos (A), WT segregant (B), *Ttgi-B3* (C), *Ttgi-A3* (D) and *Ttgi-A3/gi-B3* (E) leaf fragments in constant light (n = > 10). White bars represent the subjective day and grey bars represent the subjective night. (F) Circadian period (hours). (G) Relative Error of Amplitude (RAE). A RAE value above 0.5 (indicated by the dashed line) is considered arrhythmic. Period and RAE were calculated using Biodare2 for the CF parameter NPQ. Significant differences (P < 0.05) calculated in R using the Kruskal–Wallis test followed by post-hoc Dunn’s test.

### Circadian oscillator transcript abundance is perturbed in *Ttgi-A3/gi-B3 loss-of-function* lines

Having found that *GI* double mutants affect circadian rhythms in wheat we investigated how this might occur by examining regulation of circadian oscillator transcript abundance in *Ttgi-A3/gi-B3* in white light dark (LD) and continuous white light (LL). Under LD, transcripts of *TtLHY*, *TtTOC1*, *TtPRR73*, *TtLUX* and, *TtGI* oscillated robustly in the Kronos genotype and *Ttgi-A3/gi-B3* (Fig. 3). In LD cycles in Kronos there were robust cycles of circadian oscillator transcript abundance in both Kronos and *Ttgi-A3/gi-B3,* (Fig. 3). The loss of functional *GI* resulted in a reduction in amplitude of *TtGI* and *TtELF3*, an increase in amplitude of *TtTOC1* and little effect on the other components (Fig.3). The slightly advanced wave forms in LD in *Ttgi-A3/gi-B3* suggested that loss of *TtGI* advances the entrained phase of circadian oscillators in wheat (Fig. 3). This early phase phenotype advanced the timing of maximal expression of *TtLHY* and *TtTOC1* in *Ttgi-A3/gi-B3* compared to Kronos (*TtLHY* ZT0 compared to ZT3, *TtTOC1* ZT9-12 compared to ZT12) in LD (Fig. 3A and 3B), while phasing of peak transcript abundance of *TtPRR73* (ZT6), *TtGI* (ZT9) and *TtLUX* (ZT12) was unaffected in *Ttgi-A3/gi-B3* (Fig. 3C, 3E and 3F). In LD conditions, there were no significant differences in *TtPPD-1* peak transcript abundance between the *Ttgi-A3/gi-B3* and Kronos (Fig. 3D).

**Fig. 3.**
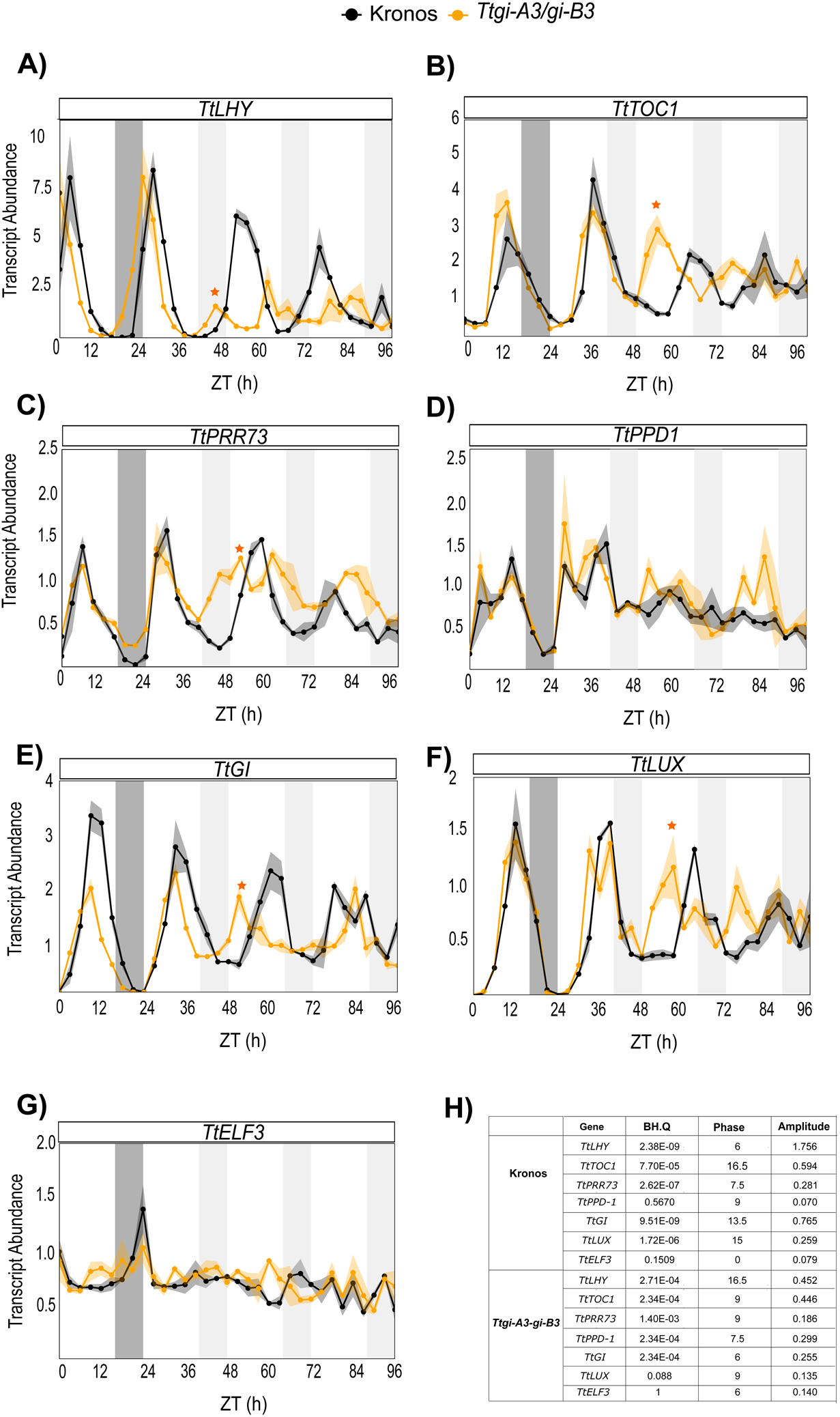
Functional *TtGI* is required for persistence of robust oscillations of circadian oscillator transcript abundance in continuous light. (A–G) \Mean abundance (± SEM, represented by the shaded ribbon) of circadian oscillator transcripts (n = 3-5) in *Ttgi-A3/gi-B3* (yellow) and Kronos WT (black). Transcript abundance (ΔΔCq) is relative to *TtRP15* and *TtRPT5A*, (A) *TtLHY*, (B) *TtTOC1*, (C) *TtPRR73*, (D) *TtPPD1*, (E) *TtELF3*, (F) *TtLUX*, and (G) *TtGI*. Orange stars indicate the peak of first true circadian cycle in LL in *Ttgi-A3/gi-B3.* White bars represent light, dark grey bars represent darkness in the last cycle before release into constant light. Light grey bars represent subjective night in constant light.

In the first true circadian cycle (beginning 24 hours after the transition to LL), peak transcript abundance of *TtLHY*, *TtTOC1*, *TtPRR73*, *TtLUX* and *TtGI* was phased earlier in *Ttgi-A3/gi-B3* than in Kronos (denoted by orange stars). Oscillations of transcript abundance were detected in Kronos in the subsequent LL cycles, albeit with reducing amplitude, whilst in *Ttgi-A3/gi-B3* transcript abundance tended to be less rhythmic (Fig. 3). Both *TtPPD-1* and *TtELF3* expression had undetectable rhythms in prolonged LL in both Kronos and *Ttgi-A3/gi-B3* (Fig. 3D and 3G), consistent with previous reports of low amplitude rhythms of wheat *ELF3* transcripts (Wittern *et al*., 2023). JTK cycle (Fig. 3.H) analysis reported that *TtGI, TtLHY, TtPPD-1* and *TtTOC1* expression as rhythmic (BH. Q < 0.05) in LL, however visual inspection suggests the rhythmic dynamics were not as robust, this was particularly evident for *TtLHY* (Fig. 3A) and *TtPRR73* (Fig. 3C).

### *TtGI* is a mild promoter of flowering in long and short-day photoperiods

Our data demonstrate that mutation of *GI* has affects wheat circadian oscillators. In Arabidopsis, *GI* contributes to photoperiodism through its role in maintaining robust circadian rhythms and through the regulation of FKF1 (Sawa *et al*., 2007). The TILLING population used in our studies carries the semi-dominant *PPD1a* allele which reduces the sensitivity of heading date to photoperiod, though the mechanism by which this occurs are not currently understood. This gave us the opportunity to investigate whether loss of *GI* function, and the associated effect on circadian oscillators can affect heading date additively to the *PPD-1*-mediated photoperiod perception pathway. We grew Kronos, WT segregant, *Ttgi-A3, Ttgi-B3* and *Ttgi-A3/gi-B3* lines in long and short photoperiods recording when each plant reached the heading growth stage (GS55, half the ear emerged from the ligule), an easily observed growth stage which is commonly used as a proxy for flowering time in wheat (Zadoks *et al*., 1974)

Under long photoperiod, heading time in the double mutant was later (59 ± 0.31 days) than either single mutant (*Ttgi-A3* 54 ± 0.78 days, *Ttgi-B3* 51 ± 0.28 days) and even later than Kronos (51± 0.51) days and WT segregant (54 ± 0.37 days) (Fig. 4A) demonstrating that the delayed flowering observed in the double mutant is caused by the absence of a functional *GI* rather than background mutations. In short photoperiods the *Ttgi-3A/gi-3B* double mutants reached GS55 10 days after *Ttgi-A3* and WT segregant, 13 days after *Ttgi-B3* and 14 days after Kronos (Fig. 4B) demonstrating an effecting on heading. These results differ from Arabidopsis where *GI* mutants (*gi-1-6*) and T-DNA insertion line (*gi-11*) flower later in long day and have minor delays in short day flowering (Fowler *et al*., 1999). Taken together these data indicate that mutation of *GI* delays flowering in both long and short photoperiods, and that mutation of either the copies of *GI* on the A or B genome alone are without effect, or very weak.

**Fig. 4.**
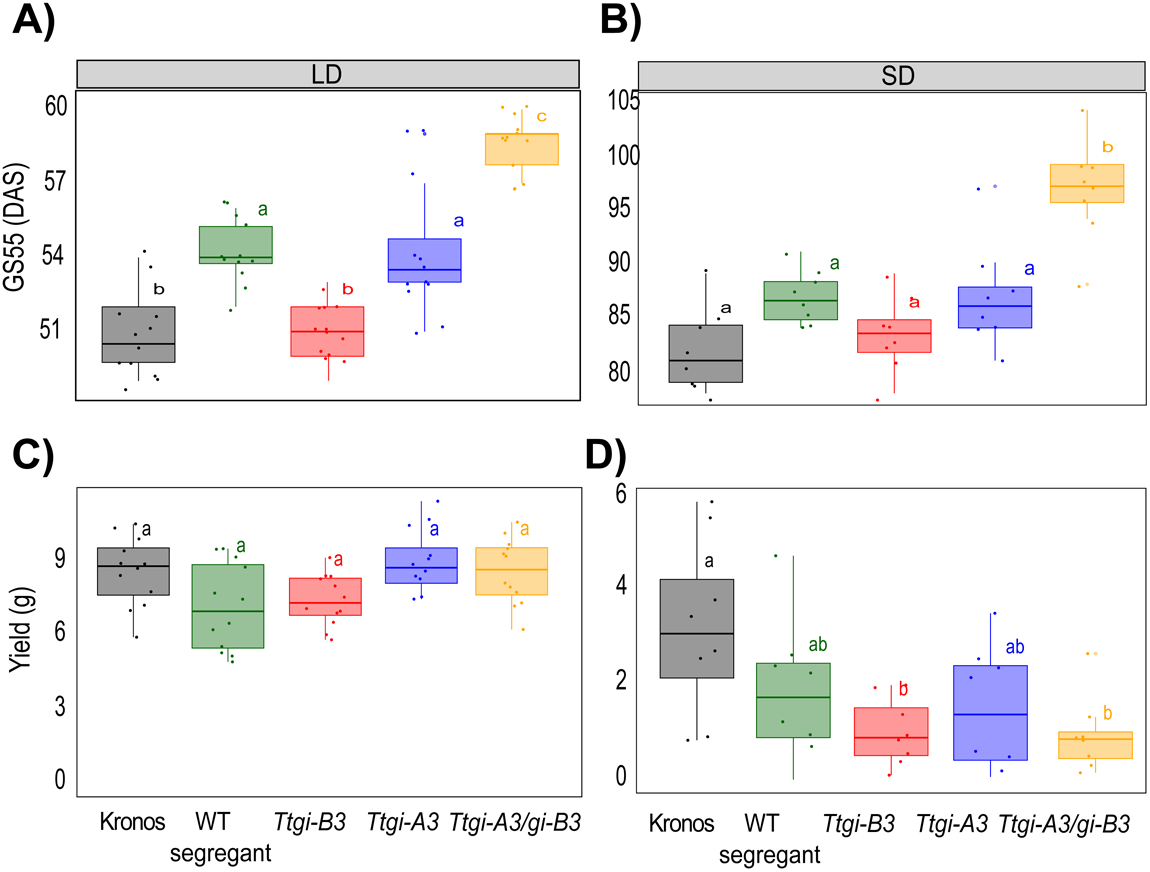
Flowering time is delayed in *Ttgi-A3/gi-B3* in long- and short-photoperiods. *Ttgi-A3/gi-B3*, *Ttgi-A3, Ttgi-B3,* WT segregant and Kronos were grown in long photoperiod (LD, 16 h light at 250 µmol m^−2^ s^−1^, 20°C: 8 h dark 16°C) and short photoperiod (LD, 8 h light at 250 µmol m^−2^ s^−1^, 20°C: 16 h dark 16°C). GS55 is used as reference for heading date and defined as days after sowing (DAS). (A) GS55 in long photoperiod (n = 12). (B) GS55 in short photoperiod (n=8). (C) Yield (total seed weight) under long photoperiod (n = 12). (D) Yield (total seed weight) under short photoperiod (n = 8). Upper and lower hinges represent the first and third quartiles (25th and 75th percentiles), the middle hinge represents the median value, whiskers represent the third quartile + 1.5*interquartile range (IQR) and the first quartile – 1.5*IQR, each dot represent individual replicates. Significant differences in heading date were tested using a Kruskal Wallis test followed by a post hoc Dunn test, and yield differences were tested using one-way ANOVA followed by Tukey’s test. The letters within each panel indicate statistical difference, samples that share the same letter in that experiment are not significantly different.

Flowering time is a key determinant of yield in wheat (Hyles *et al*., 2020). To investigate if *Ttgi-A3/gi-B3* had an effect on yield, we quantified yield traits and measured the yield of each plant (total seed weight) grown in long or short photoperiods (Fig. 4C and 4D). Predicted loss of functional *TtGI* had a greater effect on yield and yield traits under short photoperiods compared to long photoperiods. Under long photoperiods, seed yield was not significantly different between the genotypes, the mean yield per *Ttgi-A3/gi-B3* plant was 8.55 ± 0.39 g compared to 8.56 ± 0.41 g in Kronos. The mean yield of each individual Kronos plant grown in short photoperiods (3.13 ± 0.64 g) was significantly greater than *Ttgi-A3/gi-B3* (0.92 ± 0.27 g), and *Ttgi-B3* (0.99 ± 0.24 g), however there were no differences in seed weight between the WT segregant line (1.76 ± 0.30 g) and the double mutant (Fig. 4C). The mean yield of the WT segregant line was lower than in Kronos, this could be due to the presence of the mutations load in this line and that short photoperiod conditions are less favourable for optimal growth. Under long photoperiods, height, number and length of the primary head, number and weight of seeds did not significantly differ (Supplementary Fig S5). Under short photoperiod height, number of tillers and length of the primary head, were not statistically different between any of the lines, however, total seed number, seed weight and number of seeds of the primary head were substantially lower in *Ttgi-A3/gi-B3, Ttgi-A3, Ttgi-B3* compared to Kronos (Supplementary Fig S6).

### Lack of functional *TtGI* did not have large effects on heading date and yield traits in the field

We had established that *Ttgi-A3/gi-B3* had a late heading phenotype in long and short photoperiods in controlled environment conditions. Next, we wished to determine if this effect was observed under field conditions. Wheat is a commercial crop, and it was therefore important to understand if the heading date phenotype seen in the laboratory was observable in the field, where interactions with the environment are more complex.

We performed a small field trial at the NIAB experimental farm in Cambridge, UK in the 2021 field season (environmental information in Supplementary Data). Plots of Kronos, WT segregant and single A mutant were grown (*Ttgi-A3*); single B mutant was not available at the time of the trial. Five plants per plot were tagged (six plots per genotype) and the date each plant reached GS55 was recorded. *Ttgi-A3/gi-B3* reached GS55 only one day later than Kronos (*Ttgi-A3/gi-B3* 67 days, Kronos 68 days), while there was no difference in heading date between the double mutant and the WT segregant line (Fig. 5A), which could be because of the presence of background mutations within in the WT segregant affecting flowering time. Therefore, whilst Kronos is not optimised for growth in UK field conditions, we found that if *GI* affects heading in the field, the effects are modest and possibly overridden by other regulators of flowering time, particularly in a *PPD-A1a* background. The mean weight of seeds at 15% grain moisture (Kg) per head gathered from *Ttgi-A3/gi-B3* (1.45 ± 0.08) plants was not significantly different to Kronos (1.69 ± 0.08), the single A mutant (1.54 ± 0.09), or WT segregant (1.51 ± 0.07) (Fig. 5B). After the plants had fully senesced, morphological traits were measured, and a representative head was taken from each tagged plant (Supplementary Fig S7).

**Figure 5.**
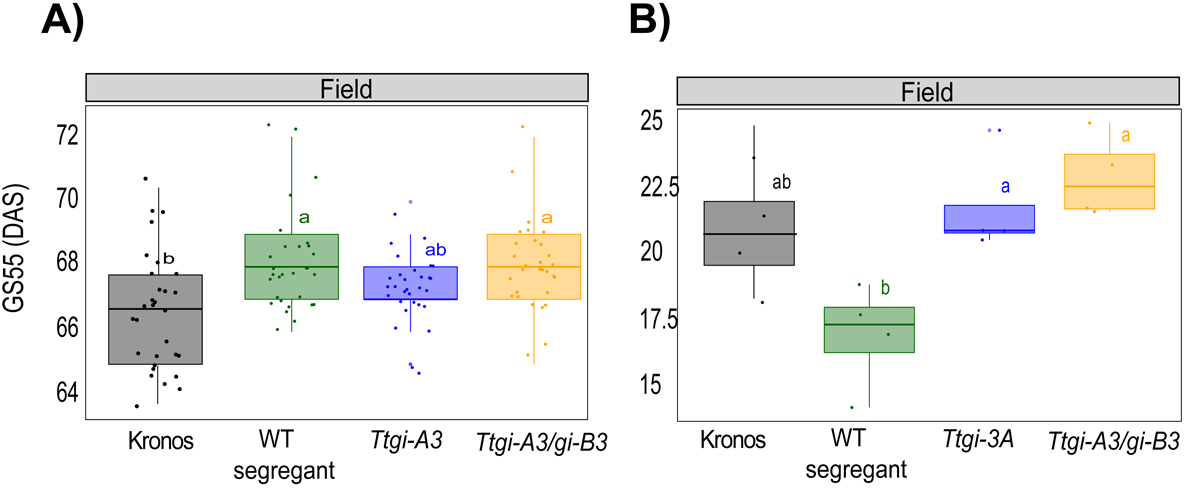
Mutations of single *TtGI* homologues and double mutant have no effect on flowering time in the field and yield. Kronos, WT segregant, *Ttgi-A3* and *Ttgi-A3/gi-B3* were grown in the field at NIAB experimental farm in the 2021 field season. (A) Heading date as defined as days after sowing to reach GS55. Data was pooled for all tagged plants of each genotype (n = 30). Significant differences in heading date were tested using a Kruskal Wallis test followed by a post hoc Dunn test. (B) Yield at 15% grain moisture in representative heads in each genotype grown in the field (n = 6). Each jitter point represents the total yield of an individual plot. Upper and lower hinges represent the first and third quartiles (25th and 75th percentiles), the middle hinge represents the median value, whiskers represent the third quartile + 1.5*interquartile range (IQR) and the first quartile – 1.5*IQR, each dot represent individual replicates. Significant differences in heading date were tested using a Kruskal Wallis test followed by a post hoc Dunn test, and yield differences were tested using one-way ANOVA followed by Tukey’s test. Significant differences in yield were tested using ONE-WAY Anova followed by Tukey’s test. The letters within each panel indicate statistical difference, samples that share the same letter in that experiment are not significantly different.

### Expression of flowering genes in the *Ttgi-A3/gi-B3* line

Having shown that *GI* is required for robust circadian rhythms, and that *TtGI* is a mild inducer of flowering in long and short photoperiods in a *PPD-A1a* background, we then characterized the effect of loss of function of *TtGI* on the transcription profile of several flowering regulatory genes. The absence of *TtGI* did not have a major effect on the expression of flowering genes at either the three-leaf stage nor GS39 (Fig. 6). Expression of *TtPPD-1* was measured using common primers for both sub-genome and sub-genome specific primers. At the three-leaf growth stage, there were two peaks of *TtPPD1* abundance at ZT 3h and 12h in both genotypes, however, the phase in *Ttgi-A3/gi-B3* was delayed for approximately 3 hours and the amplitude of transcript abundance was slightly lower than in Kronos (Fig. 6A)*. TtFT1* abundance was low in both *Ttgi-A3/gi-B3* mutant and Kronos but there was some evidence for loss of *GI* derepressing *TtFT1* at night (Fig. 6D). There were no differences in *TtCO1* and *TtCO2* expression between *Ttgi-A3/gi-B3* and Kronos (Fig. 6E-6F). Similar results were observed in growth stage GS39, *TtPPD1* amplitude was lower in the double mutant than in the Kronos line, phasing at the same time in both genotypes (Fig. 6G). No differences were detected on the expression of *TtFT1, TtCO1* and *TtCO2* between *Ttgi-A3/gi-B3* and Kronos (Fig. 6F-6H). Overall, our results show that mutation of *TtGI* does not affect the expression of photoperiodic flowering genes in wheat, contrary to Arabidopsis, indicating that there is functional diverge between the two species.

**Figure 6.**
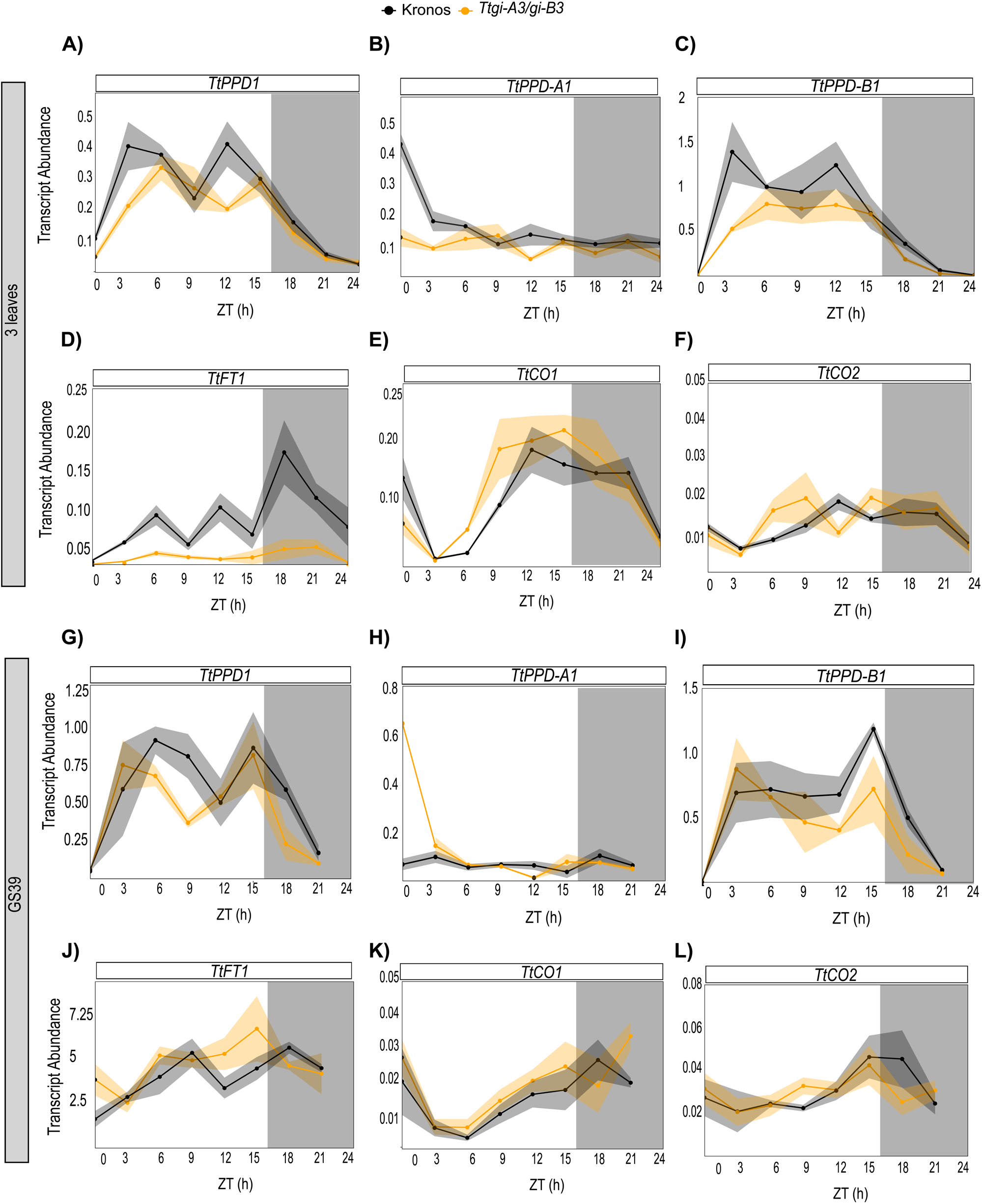
Loss of function of *TtGI* did not affect the abundance of flowering transcripts. Transcript abundance (mean ± SEM, represented by the shaded ribbon) of wheat genes involved in flowering in *Ttgi-A3/gi-B3* (yellow, n = 4-5) and Kronos WT (black, n = 4-5) grown under long day (16 h light at 250 µmol m^−2^ s^−1^, 20°C: 8 h dark 16°C). Once plants reached the 3-leaf stage or GS39, sampling of the first true leaf commenced at time 0 to 24 hours, every 3 hours. White bars represent light and grey bars represent darkness. (**A-F)** Mean abundance of flowering transcript at the 3-leaf growth stage. (**G-L)** Mean abundance of flowering transcripts at the GS39 growth stage. Transcript abundance (ΔΔCq) is relative to *TtRP15* and *TtRPT5A*, **(A-G)** *TtPPD1*, **(B-H)** *TtPPD1-A*, **(C-I)** *TtPPD1-B*, **(D-J)** *TtCO1*, **(E-K)** *TtCO2* and **(F-L)** *TtFT1*

In Arabidopsis, the phytohormone gibberellin (GA) is essential for development processes including flowering and its signalling pathway is modulated by GI (Nohales *et al*., 2019). We therefore investigated whether it was the same case in wheat by analysing the expression of two genes involved in GA biosynthesis (*TtGA20ox2* and *TtGID1*). There were no differences in expression in either 3-leaf growth stage or GS39, between the wild type and the mutant line for any of the genes analysed (Supplementary Fig. S8).

### *TtGI* sequence is conserved across modern wheat varieties

Our data indicate that whilst *GI* is required for robust circadian function, the effects on flowering in our conditions and the *PPD-A1a* background were modest. Variation within *loci* can have profound effects on the contribution of alleles to flowering, and so we investigated whether there was evidence for variation at *GI loci,* that could indicate that *GI* allelic variation might have been selected for its direct or epistatic interaction with other selected *loci* in other backgrounds and varieties. We identified *GI* orthologues in genome sequenced wheat cultivars important in modern breeding efforts (Avni *et al*., 2017; Walkowiak *et al*., 2020) (Supplementary Table S1) by performing a BLAST search using the IWGSC Ref Seq V1.1 *TaGI* homoeologues as a query sequences. We then created alignments between *GI* from the different wheat cultivars and IWGSC Refseq V1.1 to investigate variation in *GI*. Remarkably low levels of variation were observed in *GI* across the different wheat cultivars (Table 2). No variation existed in the *GI*-D coding sequences. Two SNPs within *GI*-A coding sequences were identified in the wild emmer tetraploid Zavitan, but not in any bread wheat cultivars. Five *GI*-B allele groups were identified within all varieties. The maximum number of non-synonymous SNPs in the *GI* B allele group was two. Thus, we found little variation in *GI* sequence at coding sequence level across modern wheat cultivars which can be exploited by breeders to manipulate flowering time and other circadian output traits. However, further research on non-coding sequences is needed.

**Table 2.**
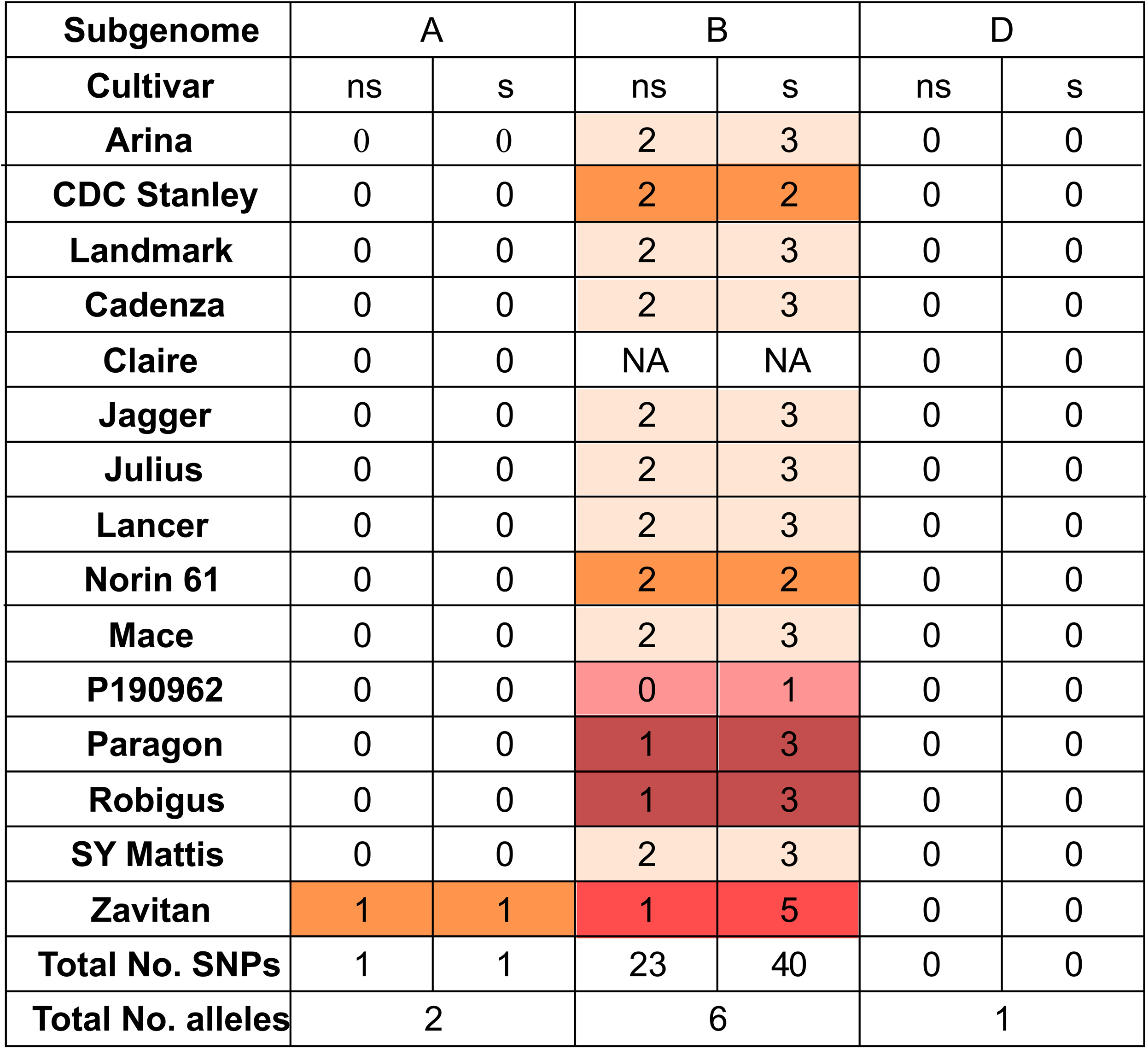
*GI* sequence is conserved across modern wheat varieties. Total number of SNPs within the CDS of *GI* across cultivars important in modern breeding efforts. A SNP was recorded if the base was different to IWGSC RefSeqv1.1. SNPs within the same codon were recorded individually. NS = non-synonymous SNP, S= synonymous SNP. Allele groups are indicated by colour. Cultivars were considered to share the same allele if the CDS were identical. Zativan is a tetraploid variety and so lacks a D subgenome. Claire *GI* B analysis was not included due to poor sequence quality.

## Discussion

### *TtGI* is required for robust circadian oscillations in wheat

Wheat circadian oscillators are perturbed in lines with predicted lack function copies of *TtGI*. *Ttgi-A3/gi-B3* lines have no detectable rhythms of NPQ or *F_v_/F_m_* in constant conditions. Similarly, circadian oscillator transcript abundance lost robust oscillations in LL. Mutation of one *GI* homoeologue (*Ttgi-A3* or *Ttgi-B3*) had no effect on the robustness of circadian rhythms in CF. This demonstrates that similar to our previous findings concerning *TtELF3*, one homoeologue of a circadian oscillator gene is sufficient to maintain robust circadian rhythms (Steed *et al*., 2021; Wittern *et al*., 2023). Due to differences in expression of circadian oscillator gene homoeologues it has been proposed that one dominant homoeologue performs the majority of the biological function in the circadian oscillator (Rees *et al*., 2022). We found that levels of transcription of both homoeologues was similar, however mutation of the *GI* A homoeologue had a greater effect on circadian period than mutation of the B *GI* homoeologue.

In Arabidopsis, GI contributes to the daily rhythms in ZTL activity (Cha *et al*., 2017; Lee *et al*., 2019). Wheat has two orthologous genes to Arabidopsis ZTL (Calixto *et al*., 2015), however it is unknown whether the two wheat ZTL orthologs are functionally redundant in wheat circadian oscillators. In Brachypodium, GI and ZTL interact in a yeast two hybrid screen (Hong *et al*., 2010). Further work is required to confirm the role of GI and ZTL in wheat at the molecular level.

### *TtGI* is a promoter of flowering in wheat

To investigate the role of *GI* in photoperiodic flowering, we measured GS55 and yield parameters in long, short photoperiod and in field conditions. Our study focuses on flowering time regulation independent of *PPD-1*-dependent pathways because the *Ttgi-A3/gi-B3* mutant was generated in a *PPD-A1a* photoperiod-insensitive background. This allowed us to investigate the role of *GI* independent of *PPD* but does means we have not been able to study genetic interactions between *GI* and *PPD-1.* In both long and short photoperiods, *Ttgi-A3/gi-B3* headed later than Kronos but few differences were observed in the field indicating that *TtGI* is a mild promoter of flowering in wheat, in a *PPD-A1a* background. In Arabidopsis, GI and FKF1 complex during the light period only during long days, promoting CO expression in a photoperiod dependent manner (Fowler *et al*., 1999; Sawa *et al*., 2007). This mechanism, plus targeting of CO for degradation by COP1 in the dark, accelerates flowering in Arabidopsis in long days and forms the molecular basis of the external coincidence model.

A core principle of the external coincidence model is that there are light sensitive and insensitive phases of the flowering promoting rhythm (Bunning, 1960). This can be demonstrated experimentally by treating plants with skeleton photoperiods (Thomas and Vince-Prue, 1997; Roden *et al*., 2002). Flowering occurred earliest in wheat plants when the light period was applied in the middle of the night (Pearce *et al*., 2017), thus demonstrating that there are rhythms in light sensitivity in the wheat flowering pathway. The circadian clock in wheat forms a complex network involving the interactions of various proteins and genes that regulations photoperiodic flowering. *ELF3* acts as a mediator between the light signalling pathway and flowering in wheat by binding to PhyB and PhyC. Moreover, in photoperiodic regulation of flowering, ELF3 regulates *PPD-1* by binding directly to its promoter and repressing its expression (Alvarez *et al*., 2023). *TtCO1/CO2* and *TtPPD1* operate in two distinct but highly connected systems, PPD-1 represses expression of *CO1* affecting the expression of *FT1* (Shaw *et al*., 2020). According to our gene expression data *TtGI* does not promote the expression of *TtCO1* and *TtCO2* under long day photoperiod which indicates that *TtGI* does not directly regulate *CO1/2*. However, a recent study in a photoperiod-sensitive line of Kronos carrying a *Ppd-A1b* allele found that *gi* mutants increase *TtCO2* expression and that CO1 and CO2 physically interact with GI in Y2H (Li *et al*., 2024). In a photoperiod-sensitive *Ppd-A1b* allele carrying line of Kronos *gi* mutants headed early with a strong interaction with photoperiod (Li *et al*., 2024). Our data support the conclusion that *GI* has little effect on heading date in a *PPD-A1a* background (Li *et al.,* 2024). Taken together these data suggest that *TtGI* regulates *TtCO1/2* through a *PPD-1*-dependent pathway and that *GI* might differentially regulate flowering in wheat and Arabidopsis.

## Conclusion

We find that reduction of *GI* function, reduces robustness of circadian oscillations but had little effect on flowering time in a *PPD-A1a* photoperiod-insensitive background. This and the recent finding that *gi* mutants affect flowering in *PPD-A1b* photoperiodic backgrounds (Li *et al*., 2024) suggest that if the circadian oscillator and *GI* do contribute to photoperiodism in wheat, it is likely to be mostly dependent on *PPD1*.

## Autor contributions

Conceptualization: Alex Webb, Mathew Hannah.

Data Curation: Laura Taylor, Gareth Steed, Lukas Wittern, Gabriela Pingarron-Cardenas

Formal analysis: Laura Taylor, Gareth Steed, Lukas Wittern, Gabriela Pingarron-Cardenas

Funding Acquisition: Alex Webb

Investigation: Alex Webb, Mathew Hannah, Laura Taylor, Gareth Steed, Lukas Wittern, Gabriela Pingarron-Cardenas

Methodology: Laura Taylor, Gareth Steed, Lukas Wittern, Gabriela Pingarron-Cardenas

Project administration: Alex Webb, Matthew Hannah

Supervision: Alex Webb

Validation: Gabriela Pingarron-Cardenas

Visualization: Laura Taylor, Gareth Steed, Lukas Wittern, Gabriela Pingarron-Cardenas

Writing – original draft: Laura Taylor, Gareth Steed, Gabriela Pingarron-Cardenas

Writing – review & editing: Alex Webb, Mathew Hannah

## Conflict of interest

Mathew Hannah is an employee of BASF. The authors declare no other competing interests.

## Funding

The work described in this manuscript were supported by UKRI BBSRC grants BB/M011194/1, BB/M015416/1, and BB/K011790/1 awarded to M.A.H. and A.A.R.W. GP-C is supported by UKRI BBSRC grant BB/W001209/1 awarded to A.A.R.W.

## Data availability

The data described in this study are available at https://doi.org/10.17863/CAM.107919

## Acknowledgements

We are indebted to Andy Greenland, Keith Gardner, Alison Bentley and other colleagues at NIAB for their support of PhD projects linked to this work. In particular, Richard Horsnell, NIAB for his assistance in growing and crossing of TILLING lines and the field team at NIAB for drilling, harvest and agronomy of *TtGI* field trial. We thank Cristobal Uauy for pre-publication access to the Kronos TILLING data and lines.

## Abbreviations

CF: Chlorophyll fluorescence
CCA1: CIRCADIAN CLOCK ASSOCIATED 1
CO: CONSTANS
LL: Continuous light
FKF1: FLAVIN BINDING KELCH REPEAT 1
FT: FLOWERING LOCUS T
GI: GIGANTEA
GS55: Growth stage 55
LD: Light dark
LHY: LONG ELONGATED HYPOCOTYL
NIAB: National Institute of Agricultural Botany
NPQ: Non-photochemical quenching
PPD-1: Photoperiod 1
PRR: PSEUDO RESPONSE REGULATOR
RAE: Relative Amplitud Error
TOC1: TIMING OF CAB EXPRESSION 1
WT segregant: Wild type
ZTL: ZEITLUPE

## Supplementary data

**Supplementary Figure S1.**
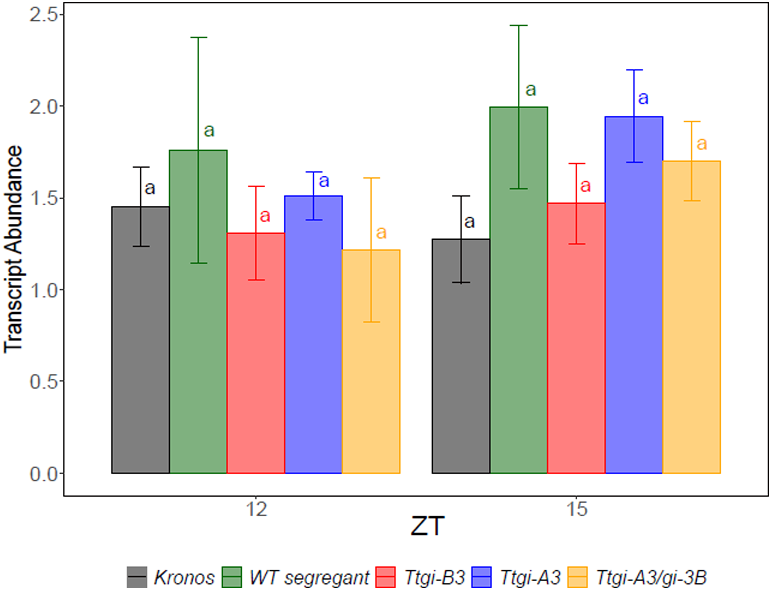
The expression of *TtGI* is consistent across both subgenomes with no significant differences between the wild type genotypes and the double mutant. *Ttgi-A3, Ttgi-B3, Ttgi-A3/gi-B3*, Kronos and WT segregant (n = 4-5) grown under long day (16 h light at 250 µmol m^−2^ s^−1^, 20°C: 8 h dark 16°C). Expression of *TtGI* was measured relative to *TtRPT5A* and *Ta22845* at ZT12 and ZT15 (mean ± SEM). The letters within the panel indicate statistical difference, samples that share the same letter in that experiment are not significantly different.

**Supplementary Figure S2.**
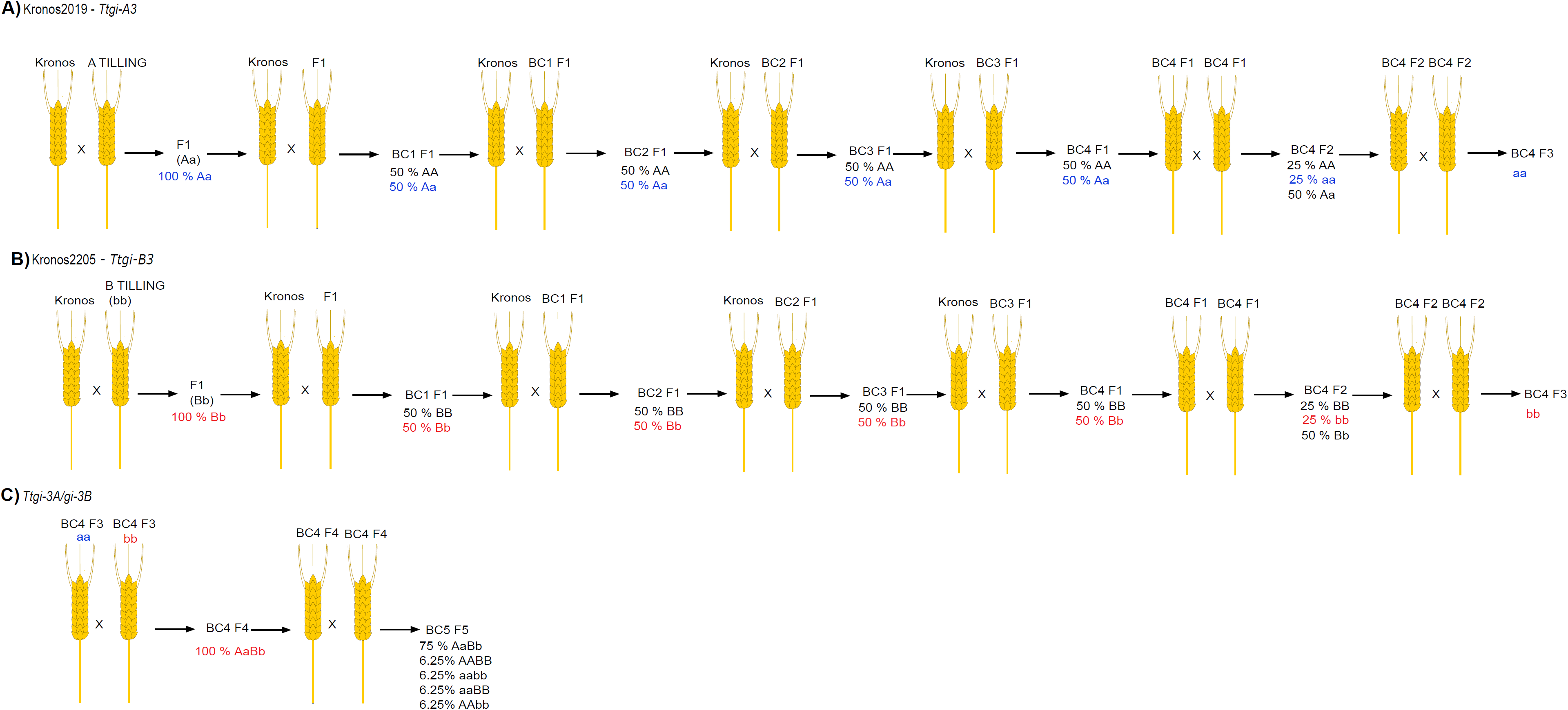
Isolation of TILLING mutants in tetraploid wheat. Crossing scheme for the creation of *Ttgi* single mutants (A), single mutant *Ttgi-A3* and (B) single mutant *Ttgi-B3.* Wild type gene is designated by a capital letter and mutated gene with lower letter. The TILLING lines *GI* Kronos2019 and Kronos2205 were crossed with Kronos background (F1). Four rounds of back crossing were completed and plants heterozygous (Aa/Bb) for the mutation selected. BC4 F1 was self-crossed to obtain BC4 F2, which was self-crossed again. Homozygous plants (aa and bb) for the mutation selected for single subgenome genotype (*Ttgi-A3* and *Ttgi-B3*, respectively) were selected. The single mutants were crossed (BC4 F4) and the progeny was self-crossed. From which the double mutants and background segregants *Ttgi-A3/gi-B3* [aabb], WT segregant [AABB]) were selected.

**Supplementary Figure S3.**
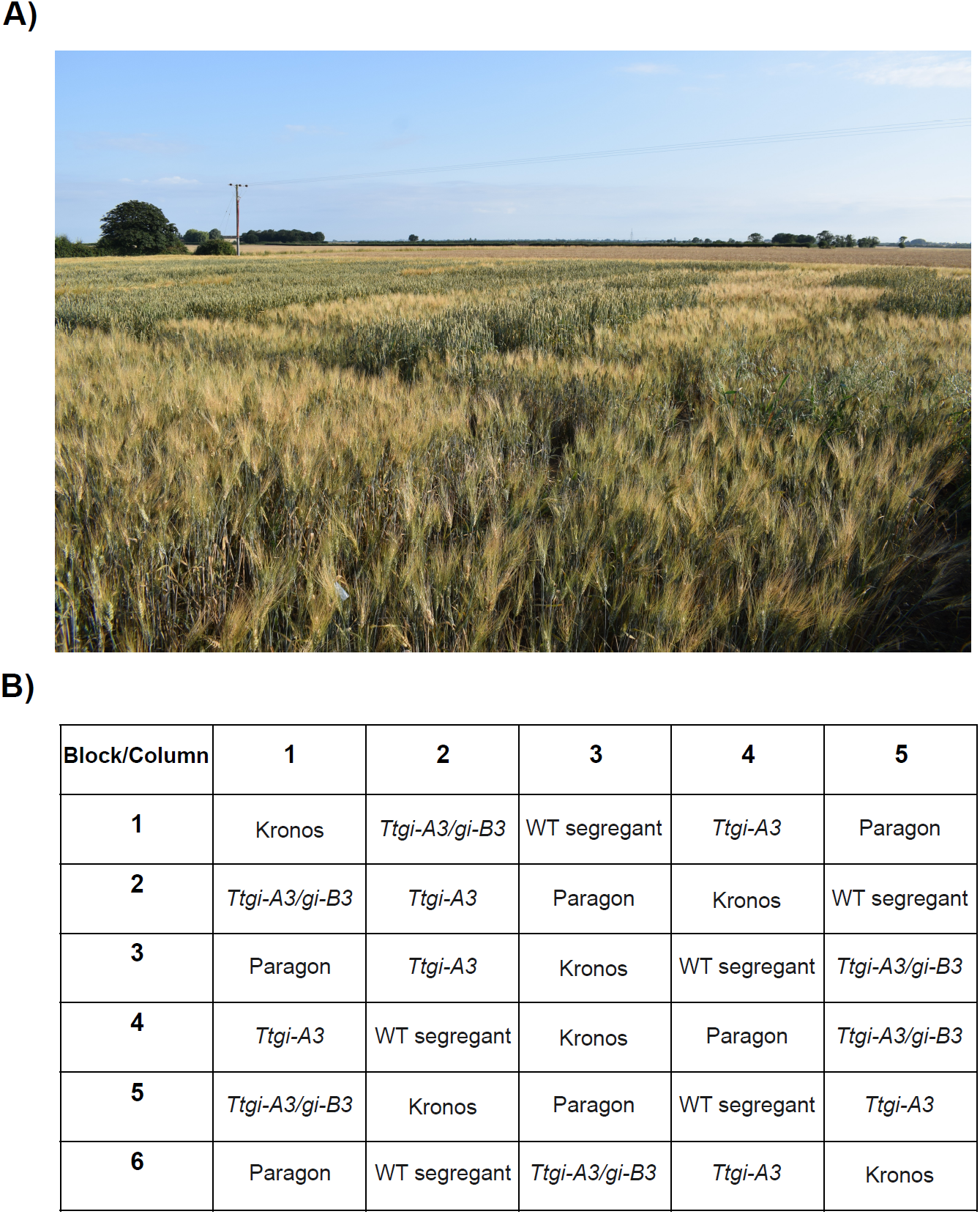
Overview of the field trial used to evaluate *TtGI* lines. A) Photograph of trial at NIAB Barr Hill field site, Cambridge, U.K). Plots of Kronos *GI* lines (awned) and Paragon controls (no awns) were flanked by Paragon plots to separate the *GI* experiment from other trials. B) Layout of trial for 2021 field season. Six plots of each genotype were drilled in a randomized block design (Paragon was not used for this analysis). Each genotype featured once in each row and a maximum of twice in each column.

**Supplementary Figure S4.**
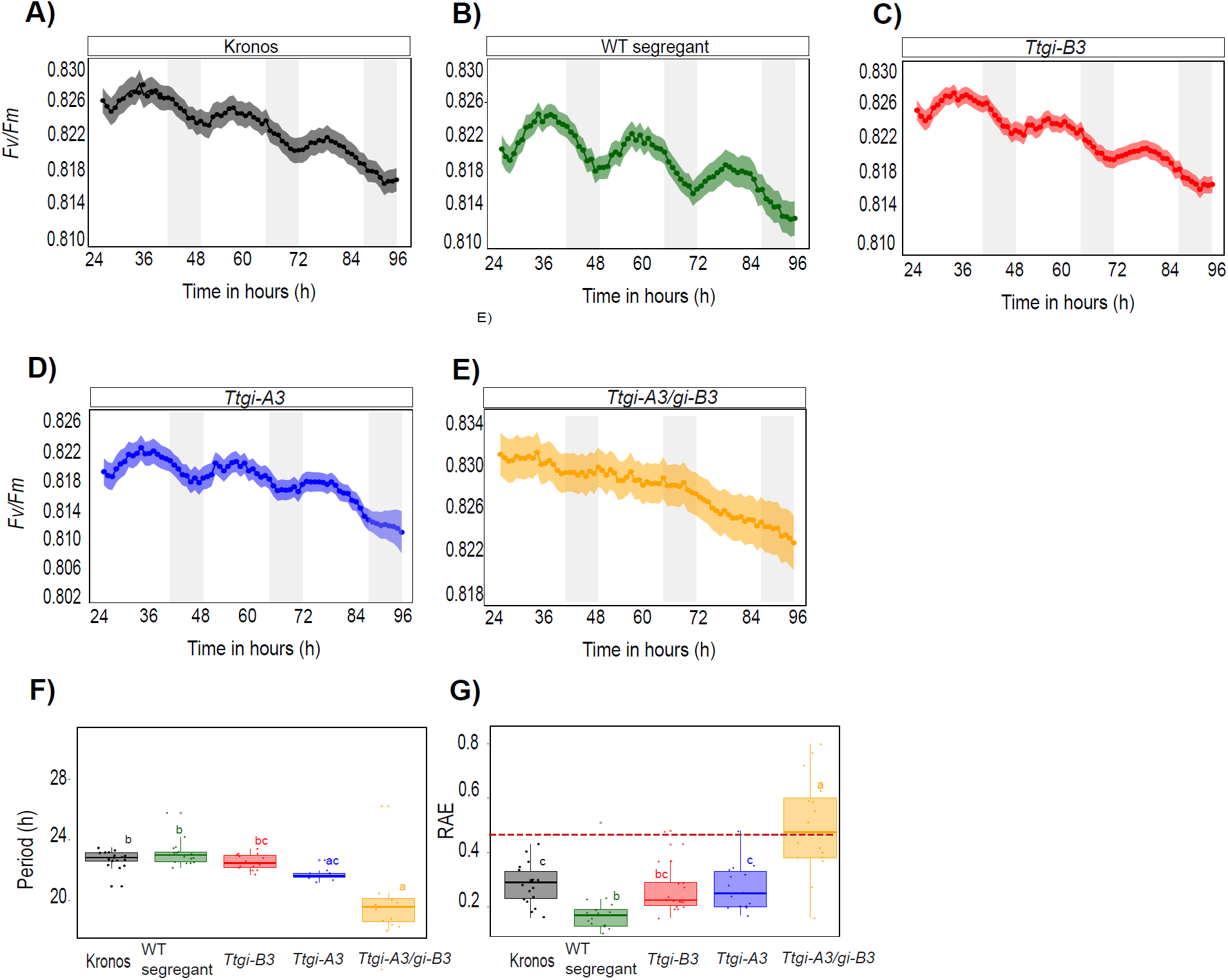
The chlorophyll *a* parameter, *Fv/Fm* is also arrhythmic in *Ttgi-A3/gi-B3* in constant conditions (light and temperature) supporting data from NPQ parameter (Figure 2). Mean of *Fv/Fm* (± SEM) of Kronos (A), WT Segregant (B), *Ttgi-B3* (C), *Ttgi-A3* (D) and *Ttgi-A3/gi-B3* (E) (n= > 10). White bars represent the subjective day and grey bars represent the subjective night. (F) Circadian period length (hours). (G) Relative Error of Amplitude (RAE). A RAE value above 0.5 (indicated by the dashed line) is considered arrhythmic. Period and RAE of *Fv/Fm* were calculated using Biodare2. Significant differences (P< 0.05) calculated in R using the Kruskal–Walli’s test followed by post-hoc Dunn’s test. (F) and (G) **t**he letters within each panel indicate statistical difference, samples that share the same letter in that experiment are not significantly different.

**Supplementary Figure S5.**
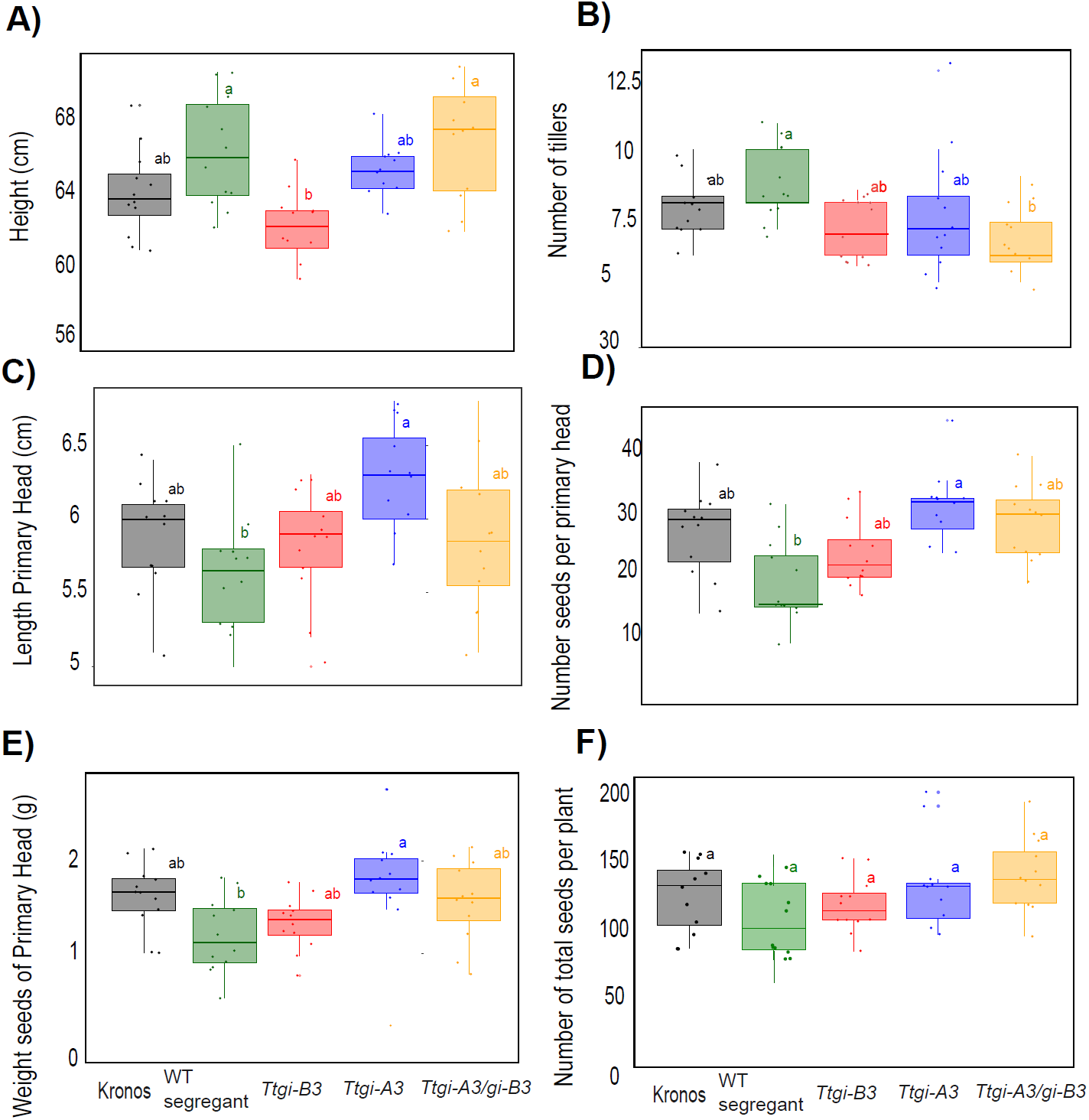
*TtGI* lines and Kronos produced equivalent yields in controlled long photoperiod. Kronos, WT segregant, *Ttgi-A3*, *Ttgi-B3* and *Ttgi-A3/gi-B3* plants were grown under long photoperiod (LD, 16 h light at 250 µmol m^−2^ s^−1^, 20°C: 8 h dark 16°C). (A) Plant length. (B) Number of tillers. (C) Length of the primary head. (D) Number of seeds per primary head. (E) Weight of seeds produced by the primary head. (F) Total number of seeds produced by each plant. Each jitter point represents an individual plant (n = 12). Significant differences were tested for using either an ANOVA with a post hoc Tukey test (plant height, primary head length, weight of seeds) or Kruskal Wallis with a post hoc Dunn test (tiller number, seed number). The letters within each panel indicate statistical difference, samples that share the same letter in that experiment are not significantly different.

**Supplementary Figure S6.**
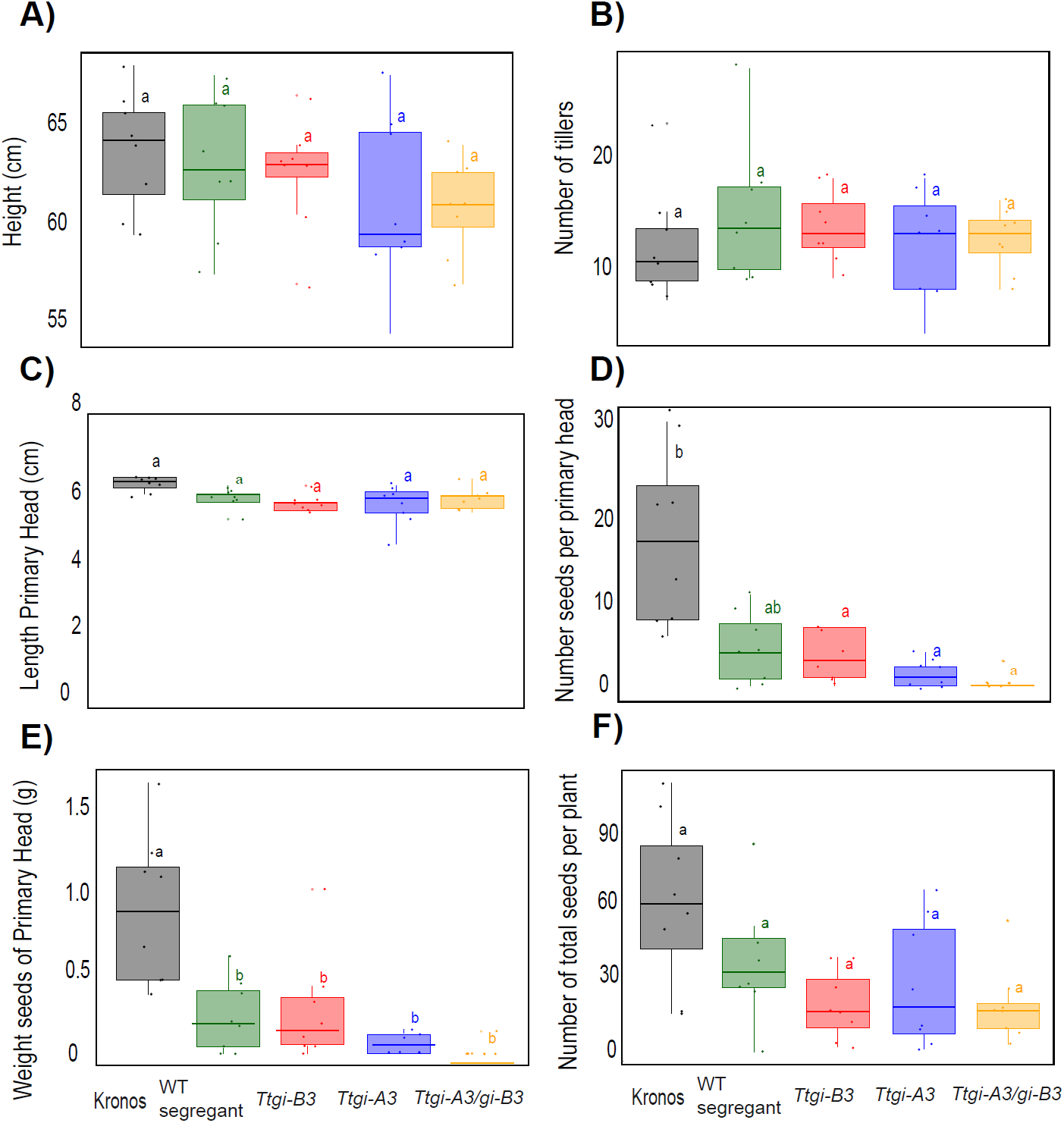
Morphological traits were broadly similar between *GI* lines and Kronos plants. Absence of a functional *TtGI* had a significantly reduced yield in controlled short photoperiod compared to Kronos. Kronos, WT segregant, *Ttgi-A3*, *Ttgi-B3* and *Ttgi-A3/gi-B3* plants were grown under short photoperiod (SD, 8 h light at 250 µmol m^−2^ s−^1^, 20°16: 16 h dark 16°C). (A) Plant length. (B) Number of tillers. (C) Length of the primary head. (D) Number of seeds per primary head. (E) Weight of seeds produced by the primary head. (F) Total number of seeds produced by each plant. Each jitter point represents an individual plant (n = 8). Significant differences were tested for using either an ANOVA with a post hoc Tukey test (plant height, primary head length, weight of seeds) or Kruskal Walli’s with a post hoc Dunn’s test (tiller number, seed number). The letters within each panel indicate statistical difference, samples that share the same letter in that experiment are not significantly different.

**Supplementary Figure S7.**
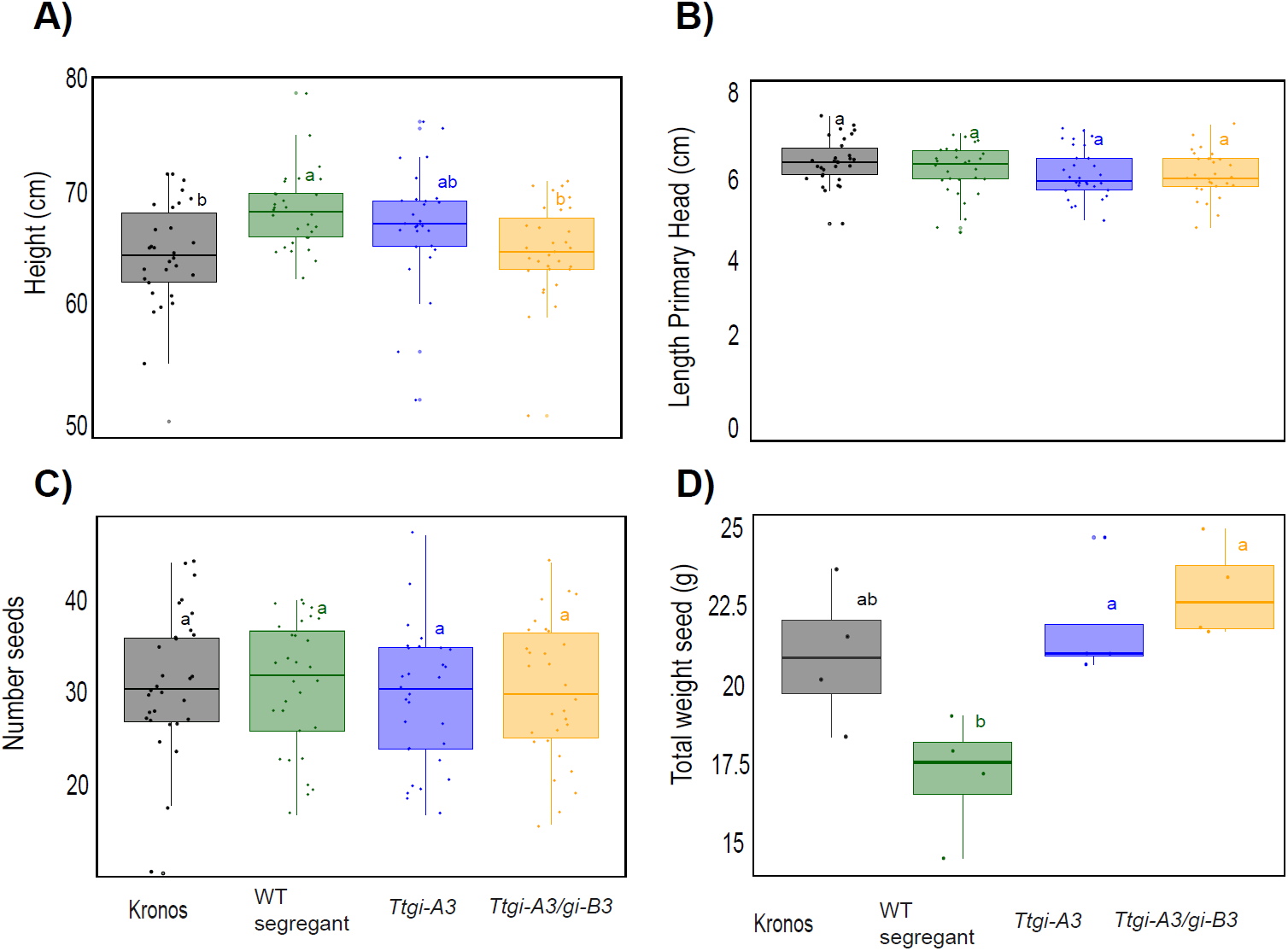
Morphological and yield traits were similar between the *TtGI* lines and Kronos in field conditions. Kronos, WT segregant, *Ttgi-A3*, *Ttgi-B3* and *Ttgi-A3/gi-B3* plants were grown in NIAB experimental farm during the 2020 field season. (A) Height. (B) Length of primary head. (C) Number of seeds of representative head. (D) Mean weight of seeds of representative head per plot. Significant differences were tested for using either an ANOVA with a post hoc Tukey test (plant height, primary head length, weight of seeds) or Kruskal Wallis with a post hoc Dunn test (seed number). The letters within each panel indicate statistical difference, samples that share the same letter in that experiment are not significantly different.

**Supplementary Figure S8.**
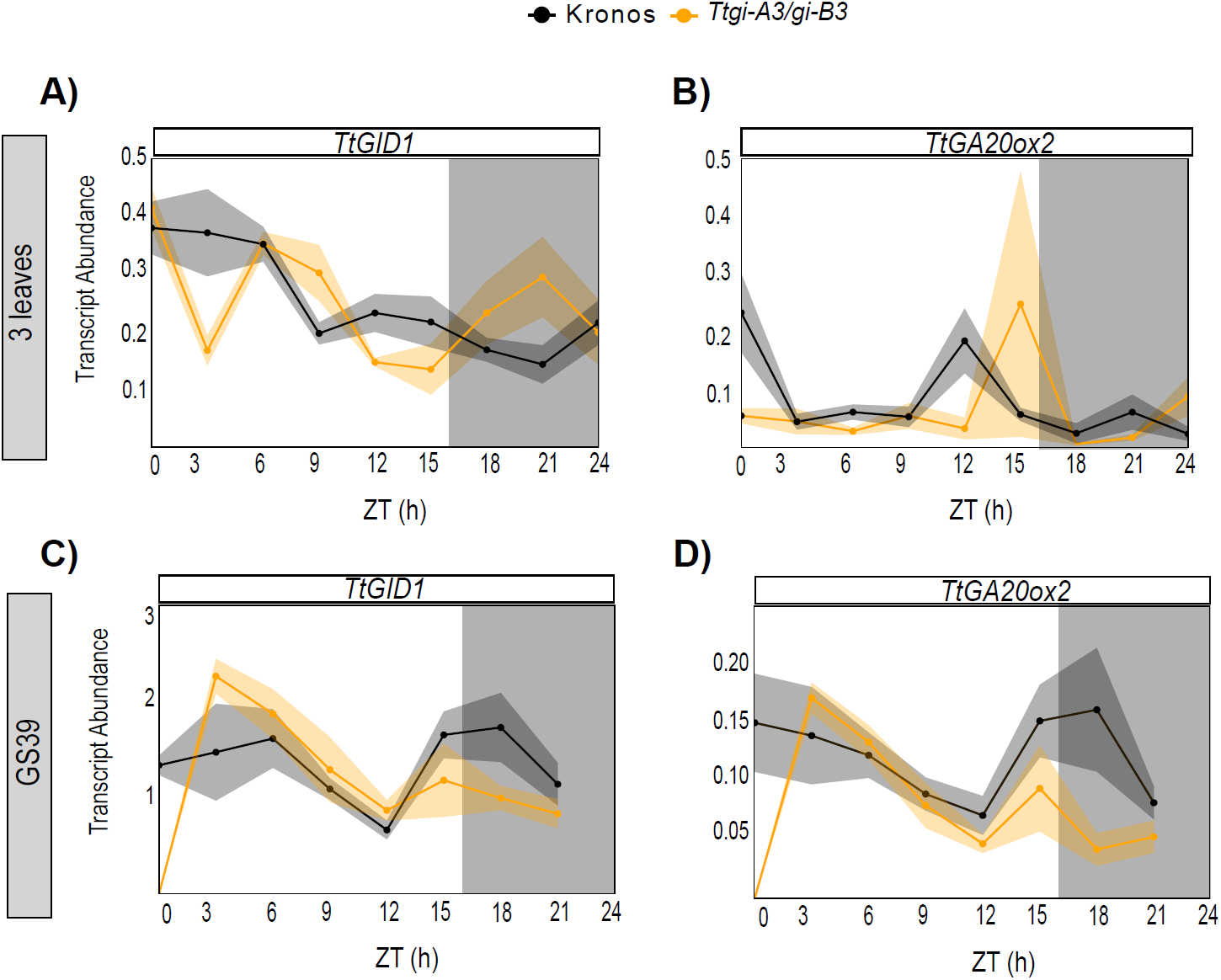
**Mutation** of function of *TtGI* did not affect the abundance of transcripts expression of genes involved in gibberellin synthesis. *Ttgi-A3/gi-B3* (yellow, n = 4-5) and Kronos WT (black, n= 4-5) grown under long day (16 h light at 250 µmol m−2 s−1, 20°C: 8 h dark 16°C). Once plants reached 3-leaves or GS39, sampling of the first true leaf commenced at time 0 to 24 hours every 3 hours. White bars represent light and black bars represent darkness. (A-F) Mean abundance (± SEM, represented by the shaded ribbon) of flowering at the 3-leaf growth stage. (G-L) Mean abundance (± SEM, represented by the shaded ribbon) of flowering at the GS39 growth stage. Transcript abundance (ΔΔCq) is relative to *TtRP15* and *TtRPT5A*, (A-C) *TtGID1*, (B-D) *TtGA20ox2*

**Supplementary Table S1.**
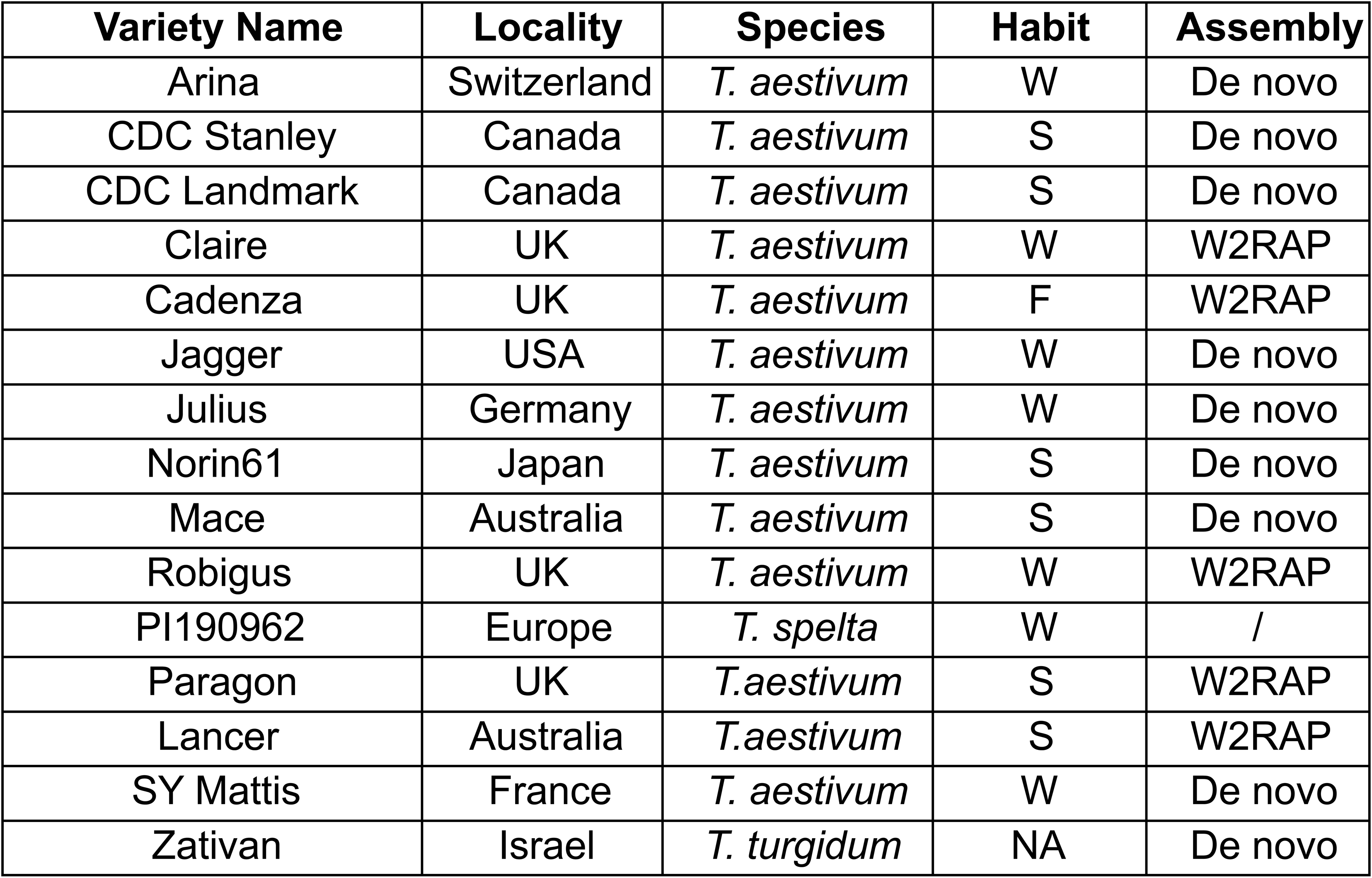
Wheat genome sequences sequenced by the wheat community (Avni et al., 2017; Walkowiak et al., 2020). Zativan (wild emmer, *T. turgidum sub spp. dicoccoides*) sequence available from Avni *et al*. 2017. All other sequences available from www.10wheatgenomes.com sequence portal and Ensembl Plants. Information for this table was gathered from www.10wheatgenomes.com and Adamski *et al*., 2020. S= spring, F= facultative and W= winter growth habit.

**Supplementary Table S2.**
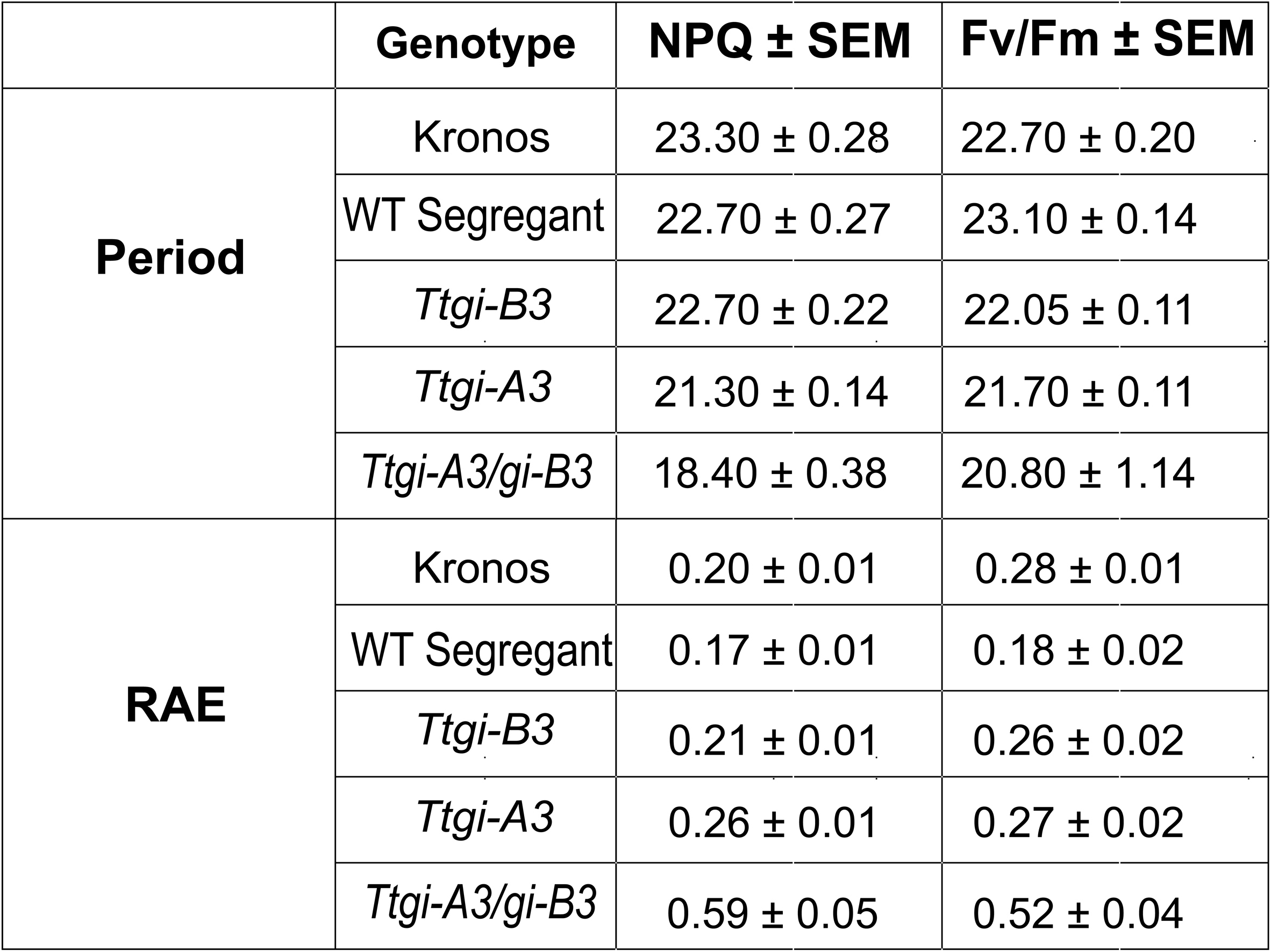
*Ttgi-A3/gi-B3* is arrhythmic for chlorophyll *a* fluorescence. *Ttgi-A3/gi-B3* is arrhythmic for chlorophyll *a* fluorescence. Period and RAE (Average ± SEM) of NPQ and *Fv/Fm* calculated using Biodare2 from Kronos, WT segregant, *Ttgi-B3*, *Ttgi-A3* and *Ttgi-A3/gi-B3* leaf fragments in constant light (n = > 10). A RAE value above 0.5 is considered arrhythmic.

## Notes

https://doi.org/10.17863/CAM.107919

### Summary of Updates

Figure 4 and associated text updated to reflect correction to error in statistical analysis for the data described in this figure; Textual changes to improve clarity in analysis of results and discussion; Supplementary information added describing the environmental conditions during field trials; Expanded analysis of circadian parameters included in revised version; Figure legends have been amended to increase clarity of description; Typographical errors corrected; Added new data and figure showing effect of mutations on GI transcript abundance; Title modified to reflect the more nuanced interpretation of the data.

